# Interleukin-11 causes alveolar type 2 cell dysfunction and prevents alveolar regeneration

**DOI:** 10.1101/2022.11.11.516109

**Authors:** Benjamin Ng, Kevin Y. Huang, Chee Jian Pua, Wei-Wen Lim, Fathima Kuthubudeen, An An Hii, Sivakumar Viswanathan, Enrico Petretto, Stuart A. Cook

## Abstract

Following lung injury, alveolar regeneration is characterized by the transformation of alveolar type 2 (AT2) cells, via a transitional KRT8+ state, into alveolar type 1 (AT1) cells. In lung disease, dysfunctional intermediate cells accumulate, AT1 cells are diminished and fibrosis occurs. Using single cell RNA sequencing datasets of human interstitial lung disease, we found that interleukin-11 (IL11) is specifically expressed in aberrant KRT8 expressing KRT5-/KRT17+ and basaloid cells. Stimulation of AT2 cells with IL11 or TGFβ1 caused EMT, induced KRT8+ and stalled AT1 differentiation, with TGFβ1 effects being IL11 dependent. In bleomycin injured mouse lung, IL11 was increased in AT2-derived KRT8+ cells and deletion of *Il11ra1* in lineage labeled AT2 cells reduced KRT8+ expression, enhanced AT1 differentiation and promoted alveolar regeneration, which was replicated in therapeutic studies using anti-IL11. These data show that IL11 maintains AT2 cells in a dysfunctional transitional state, impairs AT1 differentiation and blocks alveolar regeneration across species.

**Teaser:** Interleukin-11 stalls type 2-to-type 1 alveolar epithelial cell differentiation and prevents lung regeneration

## Introduction

The alveolar epithelium plays a pivotal role in lung homeostasis and protects the lung from inhaled environmental insults and pathogenic infections. In the alveolus, alveolar type 2 cells (AT2 cells) become activated after injury and proliferate and trans-differentiate into alveolar type 1 cells (AT1 cells) to restore alveolar structure and lung function (*1, 2*). A number of human lung pathologies, including idiopathic pulmonary fibrosis (IPF), chronic obstructive pulmonary disease (COPD) and post-infective lung damage, are characterized by failure of homeostatic AT2-to-AT1 transitions (*3, 4*).

Recent large-scale single cell RNA sequencing (scRNA-seq) studies of human pulmonary fibrosis have identified transitional cells that exhibit a dysfunctional phenotype and have a reduced capacity to differentiate into AT1 cells (*5*–*9*). These disease-associated transitional AT2 cells, coined KRT5^-^/KRT17^+^ or aberrant basaloid cells, accumulate in the lungs of patients with IPF (*6, 7*) and after severe SARS-CoV-2 infection (*10*–*12*). An analogous population of transitional cells termed KRT8+ Alveolar Differentiation Intermediate (KRT8+ ADI) / Damage-Associated Transient Progenitors (DATPS) / Pre-Alveolar type-1 Transitional cell State (PATS) are similarly seen in the damaged alveolus in mouse models of lung injury (*13*–*15*).

In mice, transitional cells, herein referred to as KRT8+ transitional cells, can be derived from either AT2 cells or airway stem cells, and possess the capacity to differentiate into mature AT1 cells (*13, 16*). Importantly, KRT8+ transitional cells in the mouse exhibit transcriptional similarities to human disease-associated KRT5^-^/KRT17^+^ / aberrant basaloid cells, including epithelial-mesenchymal transition (EMT), p53 and cell senescence pathways and expression of KRT8 itself (*13, 15*). Furthermore, KRT8+ transitional cells are thought to contribute to fibrosis via expression of profibrotic and proinflammatory mediators. Recent studies have shown that elevated TGFβ signaling in AT2 cells and IRE1α activity in DATPS maintain the KRT8+ cell state following lung injury in mice (*17*–*19*). Similarly, the persistence of senescent AT2 cells promotes progressive pulmonary fibrosis (*20*).

Interleukin-11 (IL11), a member of the IL-6 family of cytokines, is upregulated in the airways following viral infections and has been associated with a range of respiratory disorders (*21*). We previously reported that IL11 was increased in the lungs and fibroblasts from patients with IPF, and its expression correlates with disease severity (*22*). A contemporaneous study found that *IL11* was expressed in a range of cell types in fibrotic lungs of patients with Hermansky-Pudlak Syndrome (HPS) and also in *SFTPC*^+^ cells in IPF (*23*). More recent pharmacologic studies using siRNA have validated the role of IL11 in lung fibrosis (*24*).

In the current study, we leveraged single cell RNA sequencing (scRNA-seq) datasets from patients with lung disease and analyzed IL11-lineage labeled cells in a mouse model to delineate the different lung cell types expressing IL11 in disease. We examined whether IL11 signaling plays a role in alveolar regeneration via its specific activity in AT2 cells using conditional *Il11ra1* deletion in AT2 cells and lineage tracing in mice that were subjected to bleomycin lung injury and also in studies of AT2 cells in vitro. We also tested whether a neutralizing anti-IL11 antibody administered to mice with lung injury could promote alveolar regeneration by enhancing AT2-to-AT1 differentiation.

## Results

### IL11 is specifically expressed in aberrant alveolar KRT5^-^/KRT17^+^ cells in human pulmonary fibrosis

To characterize IL11 expressing cells in human pulmonary fibrosis (PF), we re-analyzed large-scale scRNA-seq data of lung cells from patients with pulmonary fibrosis from two independent studies by Habermann et. al., and Adams et. al. (GSE135893 and GSE136831 respectively) (*6, 7*). Our analysis showed that, in health, *IL11* was expressed at very low levels in the lung and its expression was barely detected across most lung cell types (**Fig S1**). In contrast, in PF, *IL11* was elevated in mesenchymal and epithelial cells populations and rarely detected in immune and endothelial cells (**Fig S1**). Within mesenchymal cells, *IL11* was most elevated in PLIN2^+^ lipofibroblasts and disease-specific HAS1^high^ fibroblasts (**Fig S1**), which supports our previous findings (*22, 25*) and further associates *IL11* with pathological fibroblast activity in PF.

Amongst the various epithelial cell types identified in the two datasets, we observed particular enrichment of *IL11* expression in disease-specific KRT5^-^/KRT17^+^ (*P*<2.2×10^−16^) and aberrant basaloid cells (*P*<2.2×10^−16^) but limited *IL11* expression in basal, ciliated, MUC5B^+^, SCGB3A2^+^, AT2, transitional AT2 or AT1 epithelial cells (**Fig 1A-D and Fig S1 and S2**). In contrast, *IL6*, which was recently implicated in airway epithelial dysfunction in fibrotic lung diseases (*26*), was broadly expressed in AT2, Mesothelial, MUC5AC+ High, MUC5B+ and Goblet epithelial cell types (**Fig S2**) but seen rarely in transitional cells in both control and PF lungs.

**Fig. 1.**
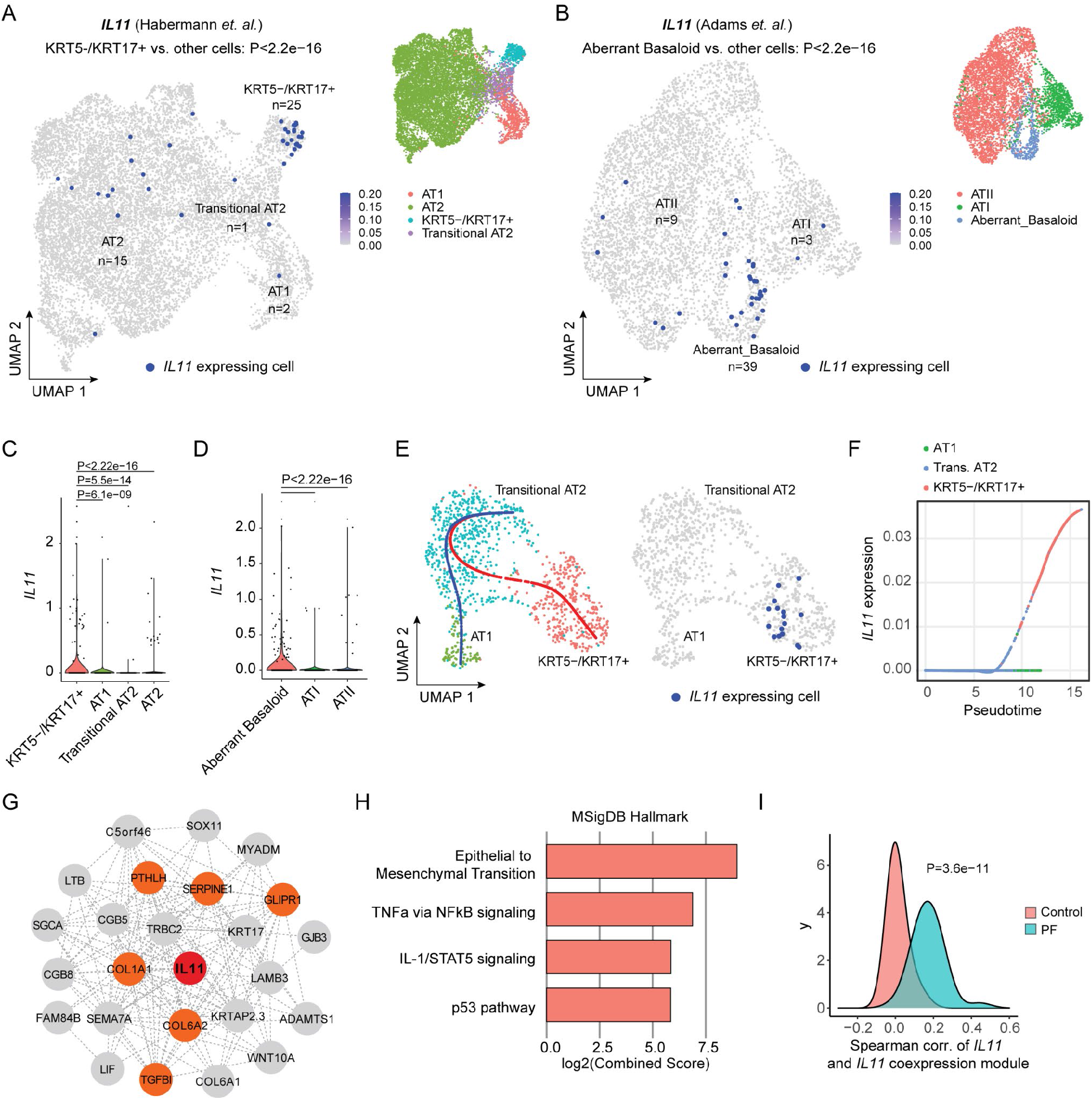
*IL11* is specifically expressed by aberrant KRT5^-^/KRT17^+^ epithelial cells in human pulmonary fibrosis. Uniform manifold approximation and projection (UMAP) visualization of *IL11* expressing single cells (colored in dark blue) in various alveolar epithelial cell populations (colored by cell types) in scRNA-seq data from Control and pulmonary fibrosis (PF) samples in (**A**) Habermann et. al. (GSE135893) and (**B**) Adams et. al. (GSE136831) datasets. The number of *IL11* expressing cells in each cell cluster is indicated. *P* values determined by hypergeometric test for enrichment in KRT5-/KRT17+ or aberrant basaloid versus other cell types. Violin plot indicating the expression of *IL11* in various alveolar epithelial cell populations in (**C**) Habermann et. al. dataset and (**D**) Adams et. al. datasets. *P* values were determined by Mann-Whitney test between KRT5^-^/KRT17^+^ or aberrant basaloid versus other cell types. (**E**) UMAP visualization of *IL11* expressing single cells colored in dark blue (right panel) and colored dots indicate cell type clustering (left panel). Blue line indicates differentiation trajectory of transitional AT2 to AT1 cells; red line indicates differentiation trajectory from transitional AT2 to KRT5^-^/KRT17^+^. Data are composed of cells from PF samples in the Habermann et. al. dataset. (**F**) Expression of *IL11* in the pseudotime trajectory from transitional AT2 to KRT5^-^/KRT17^+^ versus from transitional AT2 to AT1 cells in the Habermann et. al. dataset. (**G**) Network of genes in the *IL11* co-expression module in the transitional AT2 to KRT5^-^/KRT17^+^ cell trajectory in combined Habermann et. al. and Adams et. al. datasets. *IL11* is colored in red and genes related to epithelial to mesenchymal transition are colored in orange. (**H**) Pathway enrichment of genes in the *IL11* co-expression module using MSigDB Hallmark database and (**I**) density plot showing the spearman correlation between *IL11* and *IL11* co-expression module in Control versus PF samples in combined Habermann et. al. and Adams et. al. datasets. *P* value was determined by Kolmogorov-Smirnov test.

Since KRT5^-^/KRT17^+^ / aberrant basaloid cells may arise from defective AT2-to-AT1 differentiation, we performed trajectory and pseudotime analysis on transitional AT2 cells, KRT5^-^/KRT17^+^ / aberrant basaloid and AT1 cells on combined Habermann et. al., and Adams et. al. datasets. To do this, we first confirmed that the transcriptional profiles between aberrant basaloid and transitional AT2 and KRT5^-^/KRT17^+^ cells were highly similar (**Fig S3A**). The aberrant cells in Adams et. al. dataset were then assigned using the classification from the Habermann et. al. dataset (i.e. transitional AT2 or KRT5^-^/KRT17^+^) by Seurat’s FindTransfer Algorithm (see Methods) to obtain a consistent nomenclature across these two datasets. Our trajectory analyses showed two distinct differentiation paths for transitional AT2 cells in PF samples: 1) transitional AT2 to AT1 trajectory and 2) a trajectory from transitional AT2 to KRT5^-^/KRT17^+^ cells; with *IL11* expressed only by KRT5^-^/KRT17^+^ cells (**Fig 1E and Fig S3B**). Pseudotime analysis revealed that *IL11* was specifically upregulated along the differentiation trajectory towards KRT5^-^/KRT17^+^ cells but not towards AT1 cells (**Fig 1F and Fig S3C**).

To delineate a transcriptional program co-expressed with *IL11* along the KRT5^-^/KRT17^+^ cell trajectory, we performed co-expression analysis to the trajectory using cells assigned to the combined Habermann et. al., and Adams et. al. datasets and found that the *IL11* co-expression module was enriched for genes involved in epithelial-to-mesenchymal transition (EMT) (such as *COL1A1, SERPINE1, COL6A1, PTHLH, GLIPR1* and *TGFBI*), TNFa via NFκB signaling, IL-1/STAT5 signaling and p53 pathway (**Fig 1G-H and Table S1**). Furthermore, the association between *IL11* and the *IL11* co-expression module was highly specific to disease (**Fig 1I and Fig S3D**) suggesting a unique role of IL11 in dysfunctional alveolar epithelial cells in PF.

### IL11 is upregulated in activated AT2 cells, alveolar KRT8+ cells and stromal cells after lung injury in mice

To further characterize IL11 expressing cells in the injured lung, we used an *IL11*^*EGFP*^ reporter mouse (*27*). We performed single dose oropharyngeal injections of bleomycin (BLM), a drug that causes lung epithelial damage and fibrosis, and performed preliminary characterization of lung cells 10 days post-injury by flow cytometry. Using antibodies against a range of lung cell type markers: CD31 (endothelial cells), CD45 (hematopoietic cells), CD326 / EpCAM (epithelial cells), our analysis revealed that IL11^EGFP+^ cells were rarely observed in the uninjured lung. However, following BLM injury, we found elevated proportions of IL11^EGFP+^ cells in hematopoietic (CD45^+^ CD31^-^; *P*=.0136), epithelial (CD45^-^ CD31^-^ EpCAM^+^; *P*=.0002) and stromal cell populations (CD45^-^ CD31^-^ EpCAM^-^; *P*=.0200) (**Fig 2A-B and Fig S4**). IL11^EGFP^ was not detected in endothelial cells (CD45^-^ CD31^+^) in both injured and uninjured lungs (**Fig S4**).

**Fig. 2.**
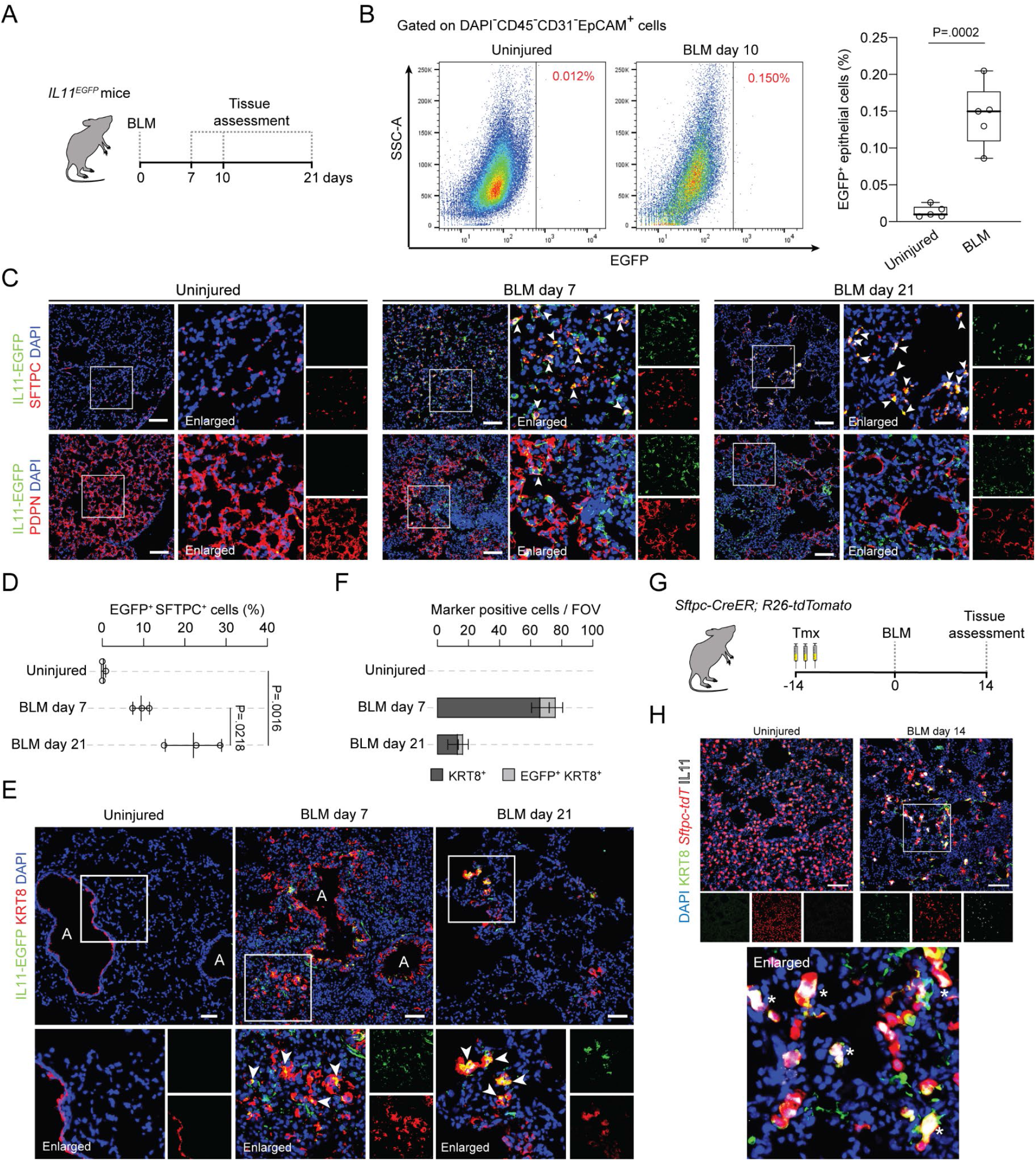
IL11 is expressed by KRT8+ transitional cells after bleomycin-induced lung injury in mice. (**A**) Schematic showing the induction of lung fibrosis via oropharyngeal injection of bleomycin (BLM) in *IL11*^*EGFP*^ reporter mice. (**B**) Representative flow cytometry analysis and proportions of EGFP^+^ CD31^-^ CD45^-^ EpCAM^+^ epithelial cells from uninjured or bleomycin (BLM)-challenged *IL11*^*EGFP*^ reporter mice. *P* value determined by two-tailed Student’s *t-*test, n = 5 mice. Representative images and quantification of immunostaining for GFP and (**C-D**) SFTPC or PDPN 7 or 21 days post-BLM challenge. Data are represented as mean ± s.d. *P* value determined by one-way ANOVA (Tukey’s test), n = 3 mice / group. (**E**) Representative images of immunostaining for GFP and KRT8 after 7 or 21 days post-BLM challenge. Airways regions are demarcated as A and white arrowheads indicate IL11^EGFP+^ KRT8^+^ cells. (**F**) Quantification of alveolar KRT8^+^ and IL11^EGFP+^ KRT8^+^ cells per field of view (FOV). (**G**) Schematic showing the period of tamoxifen (Tmx) administration and induction of lung fibrosis in *Sftpc-CreER; R26-tdTomato* (*Sftpc-tdT*) mice. (**H**) Representative images of immunostaining for KRT8 and IL11 in the lungs of *Sftpc-tdT* mice post-BLM injury. Asterisks indicate IL11^+^ KRT8^+^ *tdT*^+^ cells. Scale bars: 100 μm. DAPI for nuclei.

Since the low detection/abundance of IL11^EGFP+^ cells preclude further FACS-based analysis, we next focused on immunohistochemistry to determine the identities of IL11 expressing cells in the injured lung. To do this, we assessed the lungs of *IL11*^*EGFP*^ reporter mice at 7 or 21 days post-BLM injury by staining for GFP and counter-stained for SFTPC (AT2 marker), PDPN (AT1 marker), PDGFRA (pan-fibroblast marker) or CD45. Consistent with the flow cytometry analysis, IL11^EGFP+^ cells were very rarely observed in the lungs of uninjured *IL11*^*EGFP*^ reporter mice. In contrast, in BLM injured lungs, IL11^EGFP^ expression was notably upregulated in SFTPC^+^ AT2 cells (**Fig 2C-D**), PDGFRA^+^ fibroblasts and a subset of CD45^+^ hematopoietic cells (**Fig S5A-B**) within injured alveolar regions that were marked by areas of dense consolidation of nuclei DAPI staining. IL11^EGFP^ was localized to numerous SFTPC^+^ cells adjacent to regions of tissue disruption with an elongated morphology suggestive of AT2-to-AT1 differentiation (**Fig 2C**).

Immunostaining for an AT1 marker Podoplanin (PDPN) revealed that IL11^EGFP^ expression was very rarely detected in mature AT1 cells in injured or uninjured lungs (**Fig 2C**).

To investigate if IL11 is expressed by KRT8+ transitional cells, we performed immunostaining for GFP and KRT8 in lung sections from BLM-treated and uninjured *IL11*^*EGFP*^ reporter mice and excluded airway regions for quantification. In uninjured mice, KRT8 expression was limited to the airways whereas BLM-treatment resulted in the appearance of KRT8+ cells in the damaged alveolar regions (**Fig 2E**). There was overlap of IL11^EGFP^ expression in a proportion of KRT8 expressing cells in alveolar regions following BLM injury (**Fig 2F**).

To further test if IL11 expressing KRT8+ transitional cells are derived from AT2 cells during lung injury, we used *Sftpc-CreER; R26-tdTomato* (*Sftpc-tdT*) mice to trace AT2 cells and their descendants and monitored for the expression of IL11 specifically in this cell lineage after BLM injury. We exposed *Sftpc-tdT* mice to tamoxifen prior to BLM treatment and assessed the lungs 14 days post-injury (**Fig 2G**). We performed immunostaining using an anti-IL11 antibody, which showed consistent overlap with anti-GFP in injured *IL11*^*EGFP*^ lungs (**Fig S5C**), and counterstained for KRT8. This revealed the emergence of numerous IL11^+^ KRT8^+^ *tdT*^+^ cells with spread out/elongated morphologies 14 days after BLM injury (**Fig 2H**). IL11 and KRT8 immunostaining were not observed in alveolar regions of uninjured *Sftpc-tdT* mice, as expected. These findings revealed that IL11 expressing KRT8+ transitional cells are derived from activated AT2 cells after lung injury.

### IL11 induces cytopathic features in AT2 cells and delays AT2-to-AT1 differentiation

To investigate the functional importance of IL11 in alveolar epithelial cells, we performed 2 dimensional (2D) cultures of primary Human Pulmonary Alveolar Epithelial Cells (HPEpiC). By immunostaining, we first confirmed that HPEpiC expressed high levels of SFTPC and did not stain positive for AGER (**Fig S6A**). HPAEpiC expressed high levels of IL11RA and its co-receptor IL6ST (gp130) but lacked detectable IL6R expression (**Fig S6A**).

To test whether IL11 directly induces EMT in AT2 cells, we stimulated HPEpiC with IL11 (5 ng/ml, 24 h) and monitored for the expression of EMT related proteins (Collagen I, Fibronectin, SNAIL) along with KRT8 using immunostaining and immunofluorescence quantification (**Fig 3A**). In parallel, we treated HPEpiC with TGFβ1 (5 ng/ml; 24 h), a potent inducer of both EMT and KRT8 expression in AT2 cells (*28*–*30*), and simultaneously added a neutralizing IL11 antibody (X203) or an IgG control antibody to investigate the effect of IL11 signaling downstream of TGFβ stimulation (**Fig 3A**). This revealed that IL11 and TGFβ1 treatment led to upregulation of Collagen I, fibronectin, SNAIL and KRT8 expression, as compared to untreated cells (**Fig 3B-C**). By ELISA, we found that TGFβ1 stimulation significantly induced IL11 secretion by HPEpiC (**Fig S6B)**. The effects of TGFβ1 on the expression of EMT related proteins and KRT8 were significantly blunted by X203 (**Fig 3B-C**).

**Fig. 3.**
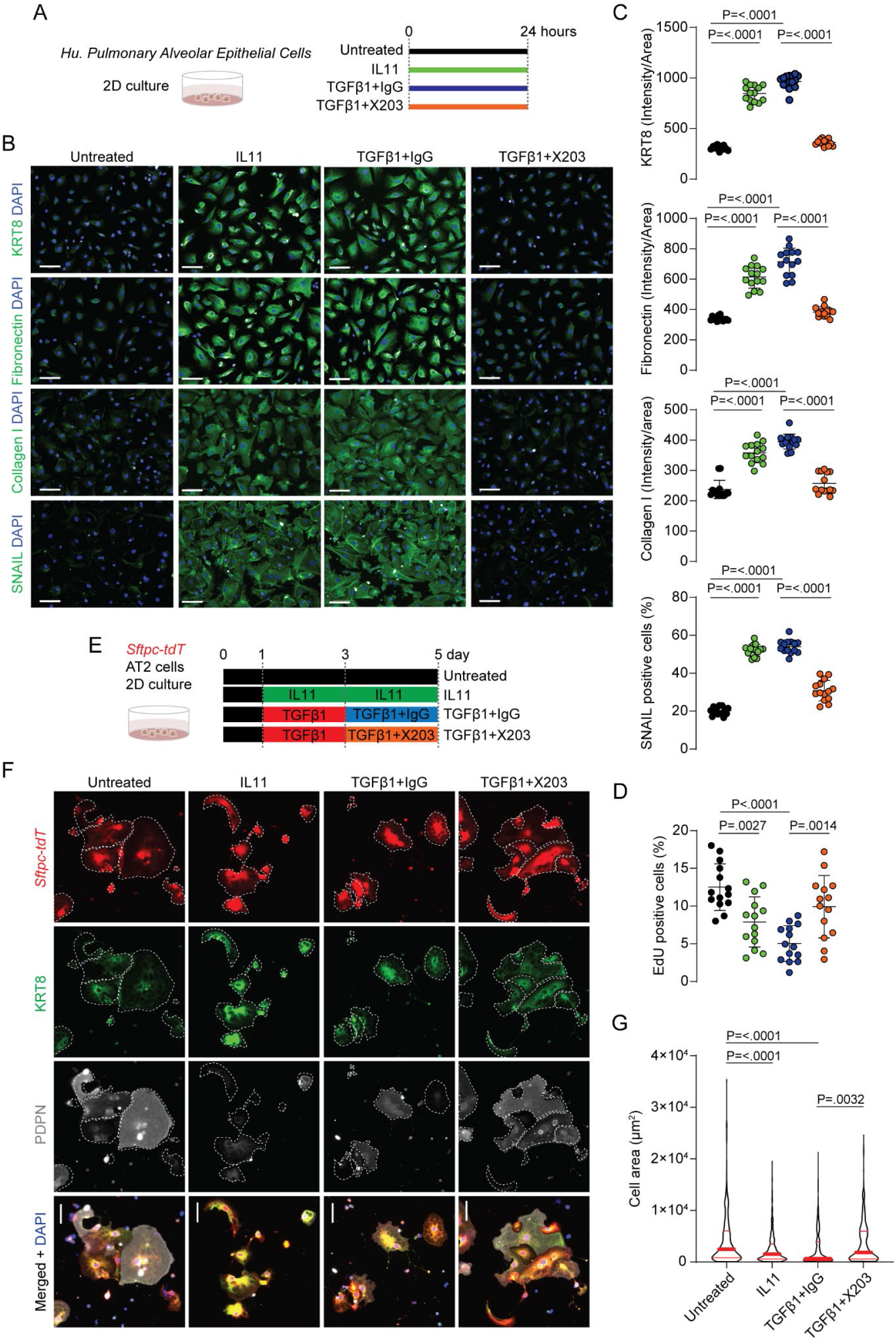
IL11 induces cytopathic features in alveolar epithelial cells and delays AT2-to-AT1 differentiation in vitro. (**A**) Experimental design for the 2D culture of primary human pulmonary alveolar epithelial cells. (**B**) Representative image of immunostaining for KRT8, Fibronectin, Collagen I and SNAIL in human pulmonary alveolar epithelial cells treated with IL11 (5 ng/ml), TGFβ1 (5 ng/ml) in the presence of anti-IL11 (X203) or IgG control antibodies (2 μg/ml). Scale bars: 100 μm. (**C**) Quantification of immunostaining in panel **B**. (**D**) Percentage of EdU positive cells in cells treated as in **A**. Data are represented as mean ± s.d. One representative dataset from three independent biological experiments is shown (14 measurements per condition per experiment). (**E**) Schematic of experimental design for the *in vitro* differentiation of AT2 to AT1 cells. (**F**) Representative images of immunostaining for KRT8, PDPN in 2D cultures of *Sftpc-tdT*^+^ AT2 cells treated with IL11 (5 ng/ml), TGFβ1 (5 ng/ml), X203 or IgG control antibodies (2 μg/ml). Scale bars: 50 μm. Individual cells are highlighted within dotted lines.(**G**) Violin plot of cell sizes of *tdT*^+^ cells. Data are pulled from two independent experiments (n > 150 cells / group). *P* values were determined by one-way ANOVA (Tukey’s test).

AT2 cell proliferation is crucial for alveolar repair after injury (*1, 31, 32*) and we tested the effects of IL11 or TGFβ1 on AT2 cell proliferation. By EdU staining, we found that exposure of cells to either IL11 or TGFβ1 (24 h) significantly impaired HPEpiC proliferation (**Fig 3D**). Furthermore, the anti-proliferative effects of TGFβ1 on HPEpiC could be reversed by X203 (**Fig 3D**). These data shows that IL11 directly induces KRT8 expression and EMT processes while impairing proliferation of human alveolar epithelial cells.

Next, we performed bulk RNA-sequencing (RNA-seq) of IL11- or TGFβ1-stimulated HPEpiC (5 ng/ml, 24 h) to evaluate the transcriptional effects of these cytokines on AT2 cells. RNA-seq analysis revealed that TGFβ1 induced transcriptomic features characteristic of KRT5^-^/KRT17^+^ cells from human fibrotic lungs (such as the elevated expression of *CDKN2A, CDKN2B, CDH2, COL1A1, FN1, SOX9, SOX4, KRT8, KRT17, KRT18* and reduced expression of *NKX2-1*) as compared to untreated cells (**Fig S6C)**. Along with these changes, *IL11* was amongst the top most upregulated genes in TGFβ1 treated HPEpiC (4.65 fold, *Padj*=1.28e-62) (**Table S2**).

In contrast to TGFβ1 treatment and consistent with data from other cell types (*22, 33*), IL11 (5 ng/ml, 24 h) did not result in significant changes in global transcription levels in HPEpiC (**Fig S6D)**, despite inducing the expression of EMT-related and KRT8 proteins (**Fig 2F-G**). Additionally, the effects of IL11 on the expression of KRT8 and EMT proteins Collagen I and fibronectin expression was blocked by the ERK inhibitor U0126 (**Fig S6E-F)**, supportive of an important role for IL11-ERK post-transcriptional gene regulation in AT2 cells. In keeping with this, we observed numerous p-ERK^+^ IL11^+^ *tdT*^+^ cells within injured regions of lungs from BLM-injured *Sftpc-tdT* mice (**Fig S6G**) indicating the activation of IL11-ERK signaling in activated AT2 cells after lung injury that was not apparent in uninjured lungs (**Fig S6G**).

AT2 cells largely increase their cell area and spontaneously undergo differentiation towards AT1 cells when cultured under prolonged 2D culture conditions. Under these conditions, AT2 cells upregulate KRT8 during early differentiation, followed by a decline of KRT8 and the subsequent upregulation of mature AT1 markers (such as PDPN) during late differentiation (*13, 29*). To test if IL11 stalls the transition of AT2 cells into mature AT1 cells, we isolated mouse AT2 cells from tamoxifen-exposed *Sftpc-tdT* mice by FACS sorting for *tdT*^+^ cells and cultured these cells under 2D culture conditions followed by treatment with IL11 (5 ng/ml) from day 1 to day 5 (**Fig 3E**). By immunostaining and cell surface area analysis of *tdT*^+^ cells, we found that numerous cells expressed PDPN and greatly increased their surface area after 5 days of culture in untreated cells (**Fig 3F-G**). In contrast, exposure to IL11 from days 1 to 5 stalled AT1 differentiation with cells expressing higher levels of KRT8, lower levels of PDPN and with reduced surface area, as compared to controls (**Fig 3F-G**).

Since prolonged TGFβ signaling impairs terminal AT1 maturation (*17, 29, 34*), we hypothesized that the maladaptive effects of TGFβ on AT1 maturation might be mediated, in part, by IL11. We tested for this by first priming AT2 cells with TGFβ1 for two consecutive days followed by subsequent treatment with TGFβ1 and X203 or IgG antibodies for an additional 2 days (**Fig 3E**). Similar to the effects of sustained IL11 treatment, we found that cells treated with TGFβ1 followed by coincubation with IgG resulted in stalled AT1 differentiation, with cells that were less elongated and expressed higher levels of KRT8 as compared to controls (**Fig 3F-G**). On the other hand, coincubation with X203 partially-relieved the stalled AT1 differentiation phenotype and significantly increased cell surface area and PDPN expression as compared to IgG-treated cells (**Fig 3F-G**). These data show that IL11 promotes AT2 cell dysfunction by causing the accumulation of KRT8+ transitional cells and delaying the terminal differentiation of AT1 cells.

### IL11 signaling in AT2 cells impairs AT2-to-AT1 cell differentiation during lung injury and promotes fibrosis

To test the hypothesis that IL11 signaling in AT2 cells promotes the accumulation of KRT8+ transitional cells and delays AT1 differentiation after lung injury in vivo, we generated AT2 cell-specific *Il11ra1* deleted mice by crossing *Sftpc-tdT* mice and *Il11ra1*^*fl/fl*^ mice to create *Sftpc-tdT; Il11ra1*^*fl/fl*^ mice. This allowed the simultaneous deletion of *Il11ra1* and the expression of *tdT* specifically in AT2 cells upon tamoxifen administration. *Sftpc-tdT; Il11ra1*^*+/+*^ mice were used as controls. We injected tamoxifen 14 days prior to BLM injury and assessed the lungs of mice 12 days post-BLM treatment (**Fig 4A**).

**Fig. 4.**
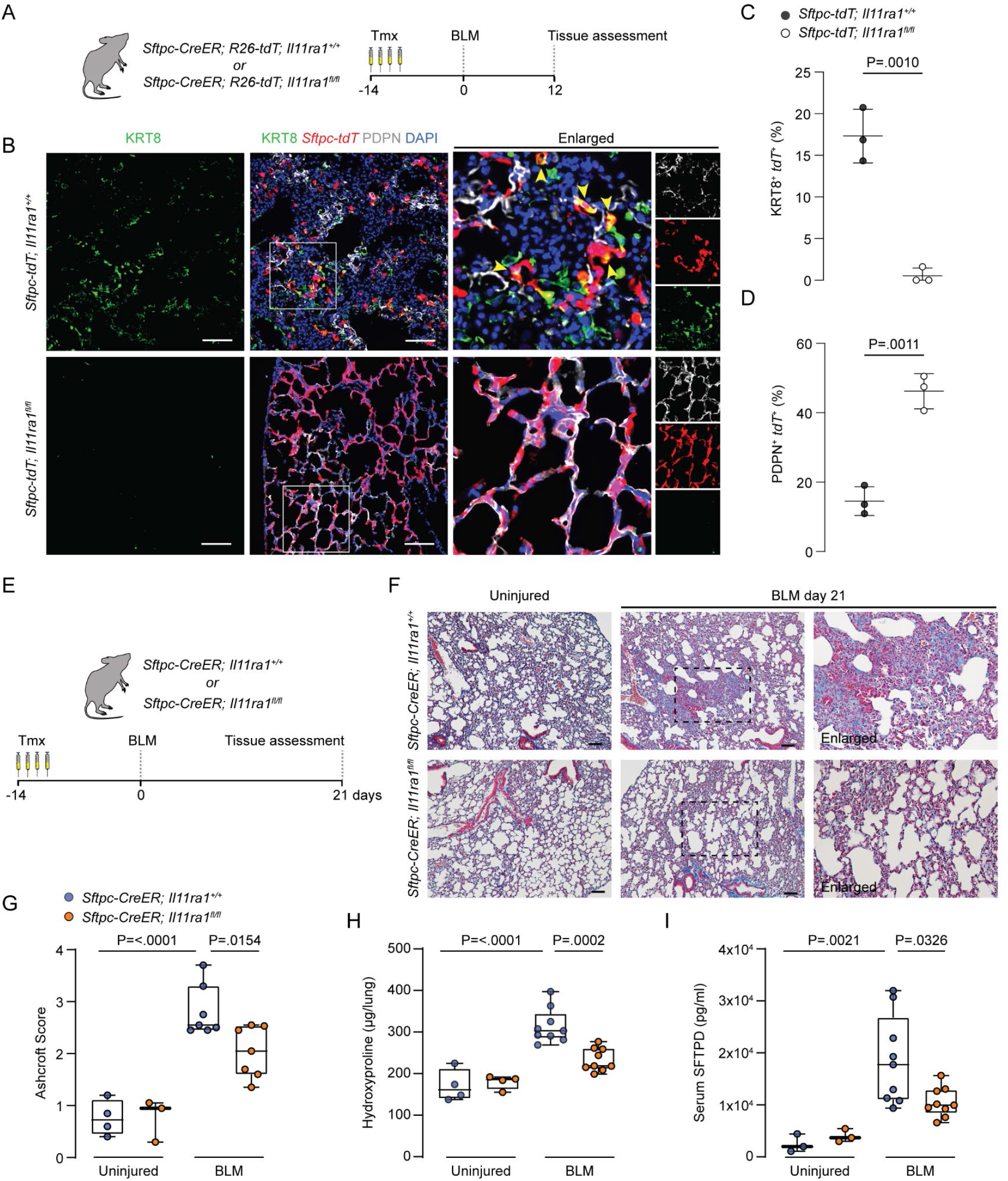
IL11 signaling in AT2 cells disrupts AT2-to-AT1 differentiation after injury and is required for lung fibrosis. (**A**) Schematic showing the induction of lung injury in *Sftpc-CreER; R26-tdT; Il11ra1*^*fl/fl*^ *or Il11ra1*^*+/+*^ mice via oropharyngeal injection of bleomycin (BLM). (**B**) Immunostaining of KRT8 and PDPN in the lungs of BLM-treated *Sftpc-CreER; R26-tdTomato; Il11ra1-floxed* or *Il11ra1*-WT mice. Scale bars: 100 μm. Yellow arrowheads indicate KRT8^+^ *tdT*^+^ cells. (**C-D**) Proportion of KRT8^+^ *tdT*^+^ or PDPN^+^ *tdT*^+^ to *tdT*^+^ cells (n =3 mice/group). *P* values were determined by two-tailed Student’s *t*-test. (**E**) Schematic showing the period of tamoxifen administration and the induction of lung fibrosis via oropharyngeal injection of BLM in *Sftpc-CreER; Il11ra1*^*fl/fl*^ *or Il11ra1*^*+/+*^ mice. (**F**) Images of Masson’s trichrome staining, (**G**) lung histopathological fibrosis scores, (**H**) lung hydroxyproline content of right caudal lobes and (**I**) serum SFTPD quantification of *Sftpc-CreER; Il11ra1*^*fl/fl*^ *or Il11ra1*^*+/+*^ mice 21 days post-BLM injury. (n = 3-9 mice / group). *P* values were determined by one-way ANOVA (Tukey’s test). Data are represented as mean ± s.d in **C** and **D** and median and whiskers extend from minimum to maximum values in **G** to **I**.

At baseline, mice with AT2 cell-specific *Il11ra1* deletion appeared histologically normal (**Fig S7A**). Following BLM-injury, immunostaining revealed numerous KRT8^+^ *tdT*^+^ cells and few newly differentiated AT1 cells (PDPN^+^ *tdT*^+^ cells) in *Sftpc-tdT; Il11ra1*^*+/+*^ control mice (**Fig 4B-D**). In contrast, parenchymal damage in BLM-treated *Sftpc-tdT; Il11ra1*^*fl/fl*^ mice was markedly reduced and KRT8^+^ *tdT*^+^ cells were rarely observed in alveolar regions from these mice.

Instead, we found numerous newly differentiated AT1 cells including regions of completely formed alveoli that were *tdT*^+^ in BLM-treated *Sftpc-tdT; Il11ra1*^*fl/fl*^ mice (**Fig 4B-D**). These results show that the deletion of *Il11ra1* in AT2 cells only, reduces KRT8^+^ cell accumulation and greatly enhances AT2-to-AT1 differentiation after BLM-injury.

To test the importance of IL11 signaling in AT2 cells for lung fibrogenesis, we used a separate cohort of *Sftpc-CreER; Il11ra1*^*fl/fl*^ mice. We administered tamoxifen 14 days prior to BLM-injury and assessed the lungs 21 days post-BLM treatment (**Fig 4E**). Tamoxifen-exposed *Sftpc-CreER; Il11ra1*^*+/+*^ mice were used as controls. The deletion of *Il11ra1* in AT2 cells from tamoxifen-exposed *Sftpc-CreER; Il11ra1*^*fl/fl*^ mice was verified by qPCR of FACS sorted CD31^-^ CD45^-^ EpCAM^+^ MHCII^+^ cells (**Fig S7B**).

Histology assessment of lungs from BLM-injured *Sftpc-CreER; Il11ra1*^*+/+*^ control mice indicated severe disruption to the lung architecture, increased collagen deposition and higher histopathological fibrosis scores, as compared to uninjured mice (**Fig 4F-G**). These pathologies were significantly reduced in mice where *Il11ra1* was deleted in AT2 cells (**Fig 4F-G**). Lung hydroxyproline content was also significantly reduced in mice lacking *Il11ra1* specifically in AT2 cells, as compared to controls (**Fig 4H**).

There was a non-significant trend of improved survival, body weights and decreased lung weights in AT2-specific *Il11ra1*-deleted mice by the end of the 21 day study period (**Fig S7C-E**). Serum surfactant protein D (SFTPD) levels, a marker of lung inflammation and epithelial injury (*35*) was elevated in BLM-injured control mice but was significantly reduced in injured mice with AT2 cell specific *Il11ra1* deletion (**Fig 4I**). Consistent with the effects seen in *Sftpc-tdT*; *Il11ra1*^*fl/fl*^ mice, immunostaining for KRT8 revealed that KRT8 expressing cells were rarely observed in the alveolar compartment of AT2 cell-specific *Il11ra1* deleted mice 21 days post-BLM injury, as compared to controls (**Fig S7F**).

### Pharmacological blockade of IL11 promotes AT2-to-AT1 cell differentiation and alveolar regeneration after bleomycin-induced lung injury in vivo

We next investigated whether anti-IL11 antibodies could promote AT2-to-AT1 differentiation and enhance alveolar regeneration when administered after lung injury. To test this, we performed BLM-induced injury to tamoxifen-exposed *Sftpc-CreER; R26-tdT* mice followed by X203 or IgG control antibody administration starting from 4 days after injury and assessed the lungs on day 12 (**Fig 5A**).

**Fig. 5.**
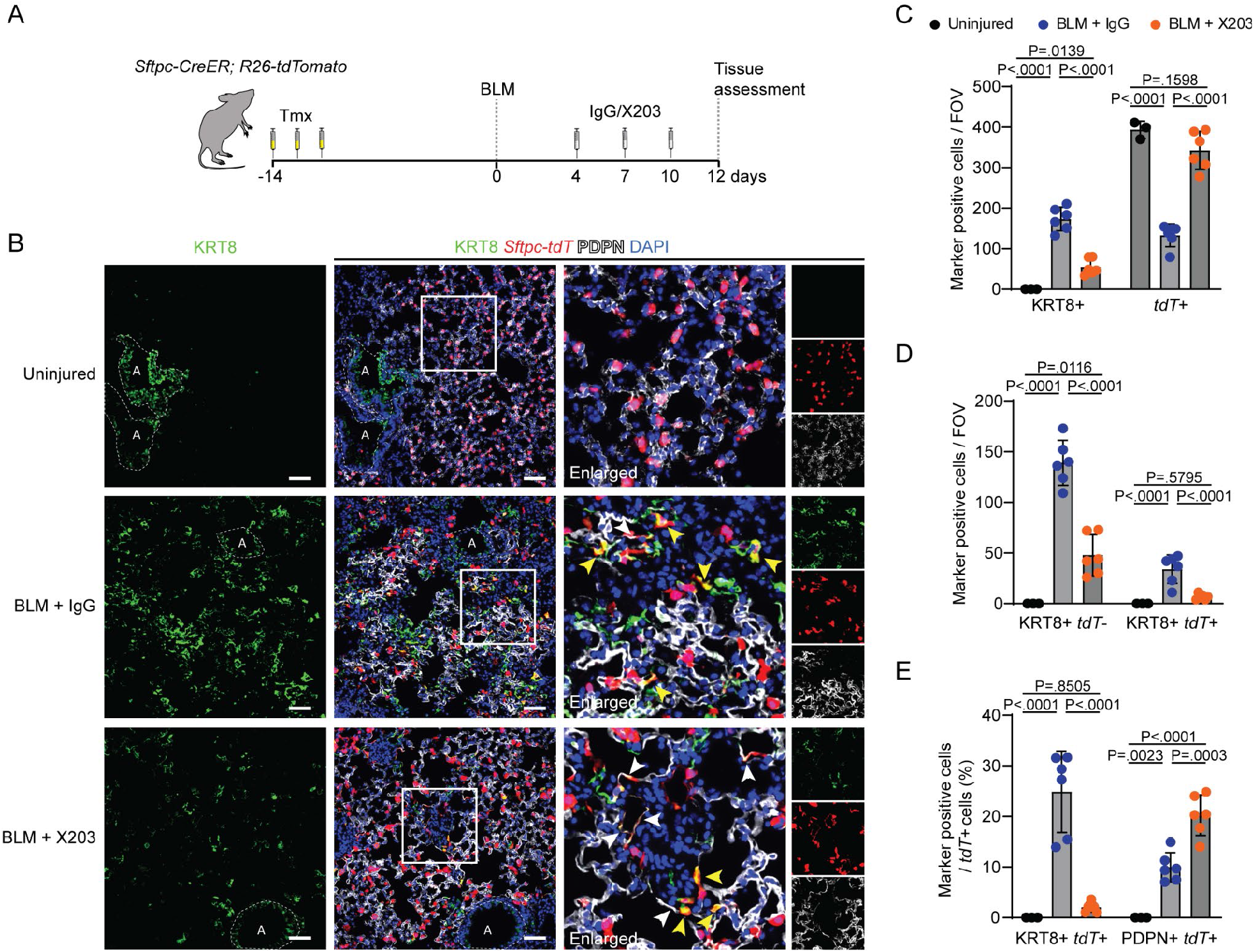
Pharmacological inhibition of IL11 enhances AT2 to AT1 cell differentiation during lung injury *in vivo*. (**A**) Schematic showing the administration time points of tamoxifen, bleomycin (BLM) and X203 or IgG antibodies in *Sftpc-CreER; R26-tdT* mice. Lung tissues were assessed at day 12 post-BLM challenge. (**B**) Representative images of immunostaining for KRT8 and PDPN in lung sections from tamoxifen-exposed *Sftpc-CreER; R26-tdT* mice treated with X203 or IgG antibodies. Yellow arrowheads indicate KRT8^+^ *tdT*^+^ cells. White arrowheads indicate flattened PDPN^+^ *tdT*^+^ cells. Recognisable airway regions are demarcated by white dotted lines. Scale bars: 100 μm. (**C**) Numbers of KRT8^+^, *tdT*^+^ and (**D**) KRT8^+^ *tdT*^+^, KRT8^+^ *tdT*^-^ per field of view (FOV), and (**E**) the proportions of KRT8^+^ *tdT*^+^ cells or PDPN^+^ *tdT*^+^ cells divided by the number of *tdT*^+^ cells in the injured lung regions from *Sftpc-CreER; R26-tdT* mice treated with X203 or IgG antibodies. n = 3 - 6 mice / group. Data are represented as mean ± s.d. *P* values were determined by one way ANOVA (Tukey’s test).

As compared to uninjured lungs, we observed widespread architectural disruption in IgG treated mice, with a large increase in KRT8^+^ cells that adopted elongated morphologies, along with a decline in the number of *tdT*^*+*^ cells (**Fig 5B-C**). Additionally, in IgG treated mice, we found an increase in non-lineage labeled KRT8^*+*^ cells (KRT8^*+*^ *tdT*^-^) that stained weakly for the AT1 marker PDPN (**Fig 5B-D**), likely reflecting an influx of airway/progenitor cells that have committed to alveolar fates in regions of severe lung injury (*13, 16, 36, 37*).

As compared to IgG treated mice, BLM-induced parenchymal damage was attenuated by X203 treatment and this coincided with reduced numbers of KRT8^*+*^ cells and proportions of lineage labeled transitional cells (KRT8^*+*^ *tdT*^*+*^ cells) and lineage negative cells (KRT8^*+*^ *tdT*^-^ cells) (**Fig 5B-E**). Furthermore, X203 treatment restored *tdT*^*+*^ cell numbers after injury to levels similar to those seen in uninjured lungs, which was associated with increased proliferation of surviving *tdT*^+^ AT2 cells as determined by immunostaining for Ki67 (**Fig 5B-C and Fig S8A**).

Consistent with the role of IL11-ERK signaling in promoting a dysfunctional KRT8 cell state in vitro, we found numerous p-ERK^+^ KRT8^+^ cells in the lungs after BLM-injury in IgG-treated mice. The occurrence of p-ERK^+^ KRT8^+^ cells was reduced in the lungs of X203-treated mice (**Fig S8B**). Finally, immunostaining for PDPN or AGER revealed that X203 treatment led to significantly enhanced differentiation of lineage labeled cells into AT1 cells (PDPN^*+*^ *tdT*^*+*^ or AGER^*+*^ *tdT*^*+*^ cells) as compared to IgG (**Fig 5B and E and Fig S8C-D**).

## Discussion

Severe respiratory diseases such as IPF and SARS-COV-2 pneumonia are associated with defects in alveolar epithelial repair and irreversible loss of alveolar epithelial cells, which ultimately leads to fibrosis and a decline in lung function. We previously discovered an important role for IL11 in lung fibrosis, mediated via its profibrotic activity in lung fibroblasts and *IL11* expression was confirmed in diseased fibroblasts in the current study (*22, 25, 38*). Here, we show that *IL11* is specifically upregulated in aberrant alveolar epithelial cells in human PF and its expression is associated with pathological pro-EMT and inflammatory gene signatures in diseased epithelial cells. In complementary studies of mice with severe lung injury, we found that IL11 is expressed by activated AT2 cells and KRT8+ transitional cells.

Due to the complex signaling milieu that occurs following severe lung injury, multiple pathways likely contribute to the emergence and maintenance of KRT8+ transitional cells, among which TGFβ, which shows IL11 dependency for its effects, is of particular importance (*17, 34*). While inflammatory cytokines such as IL-1β and TNFα induce AT2 cell proliferation, IL-1β also primes a subset of *Il1r1*-expressing AT2 cells for differentiation into DATPS (*14, 39*). Intriguingly, while IL6 is a therapeutic target in some forms of PF (*26, 40*), we show that the cell types expressing IL6 in the fibrotic lung differ from those expressing IL11 and only IL11 expression is enriched in aberrant epithelial cells.

Our data identify an IL11-stimulated ERK-dependent signaling pathway that promotes and maintains AT2 cells in a KRT8+ state and induces EMT features in AT2 cells. Furthermore, we found that the anti-proliferative effects of TGFβ and its induction of EMT genes and KRT8 expression in AT2 cells was, in part, mediated by IL11 signaling. These findings may have implications for other airway/lung disorders such as Hermansky-Pudlak Syndrome-associated pulmonary fibrosis, severe asthma and severe viral pneumonitis, including SARS-COV-2 infection, where elevated IL11 levels are elevated and implicated in disease pathogenesis (*11, 23, 41*–*43*).

Specialized lung mesenchymal cells form a supportive niche that maintains the progenitor properties of AT2 cells under homeostatic conditions (*44, 45*). In disease, impaired alveolar repair may arise due to disruption of this supportive niche and the development instead of a profibrotic niche composed of pathological fibroblasts and dysfunctional alveolar epithelial cells (*46*). Given the elevated expression of IL11 in aberrant mesenchymal and epithelial cell types in PF and its role in both fibroblasts activation and AT2 cell dysfunction, we propose that IL11 may cause multiple aspects of pathobiology across cell types in the diseased niche.

There are limitations to our study. Although several recent studies have shown that IL11 is upregulated in the lungs and *SFTPC*^+^ cells from patients with IPF (*22*–*24*), in situ studies of IL11 expression in diseased human lung tissue are required to further validate findings. We did not dissect the specific cell type expressing IL11 that impacts AT2-to-AT1 differentiation, although our earlier studies suggest a dominant role for IL11 secretion from fibroblasts for fibrosis phenotypes (*25*). In light of recent evidence highlighting the importance of distal airway secretory/basal cells in aberrant alveolar repair and fibrosis (*46*–*49*), the effects of IL11 on the recruitment and differentiation of airway / progenitor cells towards KRT8+ and AT1 cells requires study.

In conclusion, we suggest that IL11 causes lung pathology in severe lung disease through at least two pathological processes. First, causing AT2 dysfunction and maintenance of a KRT8+ cell state, thus limiting terminal AT1 differentiation and impairing alveolar regeneration. And second, stimulating fibroblast-to-myofibroblast transformation that leads to lung fibrosis and inflammation (*25*). Hence, anti-IL11 therapeutics, which are advancing towards clinical trials in patients with PF, may promote alveolar regeneration and mitigate lung fibrosis that would differentiate anti-IL11 therapy from anti-fibrotics currently used in the clinic.

## Materials and Methods

### Computational analysis of scRNA-seq datasets of human pulmonary fibrosis

Processed human IPF scRNAseq dataset by Habermann et al and Adams et. al., were downloaded from GEO with the accession number GSE135893 and GSE136831 respectively. Cell-type annotations and Uniform Manifold Approximation and Projection (UMAP) coordinates provided by the authors were used in subsequent analyses.

#### Trajectory analysis

We re-classified alveolar epithelial cells in the Adams et. al., dataset with cell-type annotations defined by Habermann et. al., using Seurat’s default label transfer pipeline. Quality of label transfer was evaluated by Jaccard Index (See *Assessment of transcriptomic similarities between epithelial cell-types* below). Transitional AT2, KRT5-/KRT17+, and AT1 cells were extracted from Habermann and Adams et al dataset for Slingshot trajectory analysis (Slingshot 1.4.0) (*50*), and the analysis was performed separately for each dataset. Briefly, Slingshot derives differentiation paths from a specified origin and calculates for each cell a pseudotime, which approximates differentiation progression of a cell toward the destination of the trajectory. In this analysis, transitional AT2 cells were specified as the origin, and two differentiation trajectories were derived, one to KRT5-/KRT17+ cells and the other to AT1 cells. Change in IL11 expression was evaluated along the two trajectories by fitting a Generalized Additive Model (GAM) with the expression of IL11 against pseudotime.

#### Assessment of transcriptomic similarities between epithelial cell-types

We examined transcriptional similarities of different epithelial clusters using the Jaccard index (a cluster here refers to cells of the same cell-type from a specific study, e.g., AT2 cells from Habermann et al dataset). First, we performed differentially expressed gene (DEGs) analysis in epithelial cells from the same study and for each cell-type retained upregulated DE genes with log2 fold change (log2FC) above the 85th percentile of the FC distribution and discarded genes with expression proportion in a cluster less than 40% compared to other cell-types. We refer to these genes as “markers’’ of a cluster, and a Jaccard index value was derived for all possible cluster pairing (of all epithelial clusters pooling together both datasets). A Jaccard index between a cluster A and a cluster B was calculated by dividing the size of the intersection of their markers over the size of the union of their markers.

#### Network analysis

Cells assigned to the differentiation trajectory from transitional AT2 to KRT5-/KRT17+ cells by Slingshot analysis were selected for IL11 co-expression analysis, done individually in Habermann et. al., and Adams et. al., dataset. Briefly, spearman correlations were calculated between the expression of IL11 and genes expressed in the selected cells. Genes with spearman correlation with FDR < .2 were kept. In summary, 103 genes were found to be significantly correlated with IL11 in Adams et al dataset, 378 genes in Habermann et al dataset, and 32 genes in both datasets. Using the R package EnrichR (enrichR 2.1.0) (*50, 51*), functional pathway enrichment analysis was performed on genes significantly correlated with IL11 (in individual datasets and combined) querying several annotation database including KEGG 2019 and MSigDB Hallmark 2020. Pathway terms with FDR < .1 were retained. De-novo network construction was performed on the 32 genes significantly correlated with IL11 in both datasets. Each node in the network represents a gene and each edge (connecting a pair of genes) the spearman correlation between the expression of the two genes in transitional AT2 and KRT5-/KRT17+ cells from Habermann et. al., dataset. Graphical representation of the network was constructed in Cytoscape (Cytoscape 3.8.2) (*52*), and genes overlapping with MSigDB Hallmark EMT process were colored.

### Mouse studies

The animal experiments were approved and conducted in accordance with guidelines set by the Institutional Animal Care and Use Committee at SingHealth (Singapore) and the SingHealth Institutional Biosafety Committee. Animals were maintained in a specific pathogen-free environment and had ad libitum access to food and water. The following mice strains were maintained on C57BL/6 background and used for the study: Sftpc^tm1(cre/ERT2)Blh^ (*Sftpc-CreER*) (*53*), B6.Cg-*Gt(ROSA)26Sor*^*tm9(CAG-tdTomato)Hze*^/J (*R26-tdTomato* mice), C57BL/6-*Il11ra1*^*em1Cook*^/J (*Il11ra1*^*fl/fl*^ mice) (*25*), C57BL/6-*Il11*^*tm1*.*1Cook*^/J (*IL11*^*EGFP*^ reporter mice) (*27*). *Sftpc-CreER* mice were crossed with *R26-tdTomato* mice to generate *Sftpc-CreER; R26-tdTomato* (*Sftpc-tdT*) mice for lineage tracing experiments. To model the deletion of *Il11ra1* in AT2 cells, *Sftpc-CreER* mice were crossed with *Il11ra1*^*fl/fl*^ mice to generate *Sftpc-CreER; Il11ra1*^*fl/fl*^ mice. *Sftpc-tdT* were further crossed with *Il11ra1*^*fl/fl*^ mice to generate S*ftpc-CreER; R26-tdTomato; Il11ra1*^*fl/fl*^ mice. *Sftpc-tdT* mice were injected intraperitoneally with 3 doses of 100 mg/kg tamoxifen (Sigma-Aldrich) starting from 14 days prior to bleomycin administration. *Sftpc-CreER; Il11ra1*^*fl/fl*^ mice and *Sftpc-CreER; R26-tdTomato; Il11ra1*^*fl/fl*^ mice were injected intraperitoneally with 4 doses of 75 mg/kg tamoxifen (Sigma-Aldrich) starting from 14 days prior to bleomycin administration. Therapeutic doses of monoclonal anti-IL11 (X203, Aldevron) were established previously (*22*). X203 or IgG control antibodies were injected intraperitoneally at 20 mg/kg starting from day 4 and subsequently on day 7 and day 10 post-bleomycin administration.

### Bleomycin model of lung injury

The bleomycin model of lung fibrosis was performed as previously described (*22*). Briefly, female mice at 10-14 weeks of age were anesthetized by isoflurane inhalation and subsequently administered a single dose of bleomycin (Sigma-Aldrich) oropharyngeally at 0.75 U/kg body weight (for *IL11*^*EGFP*^ reporter mice) or 1.5 U/kg body weight (for all other mouse strains) in a volume of saline not exceeding 50 μl per mouse. Uninjured control mice received equal volumes of saline oropharyngeally. Mice were sacrificed at indicated time points post-bleomycin administration and lungs were collected for downstream analysis.

### Mouse lung dissociation and FACS analysis

Mice lung dissociation was performed as previously described with slight adjustments (*54*). Briefly, lungs were perfused with cold sterile saline through the right ventricle. The lungs were then intratracheally inflated with 1.5 ml of Dispase 50 U/ml (Corning) followed by installation of 0.5 ml of 1% low melting agarose (BioRad) via the trachea. The lungs were excised and incubated on an orbital shaker for 45 min at room temperature. Each lobe was then minced into small pieces in DMEM (GIBCO) supplemented with 10% FBS (GIBCO) and 0.33 U/ml DNase I (Roche) and placed on the orbital shaker for an additional 10 minutes. The cells were then filtered through a 100 μm cell strainer and centrifuged at 400 g for 5 min at 4°C. The cell pellet was resuspended in ACK-buffer (GIBCO), incubated for 2 min on ice and then filtered through a 40 μm cell strainer. The cells were centrifuged at 400 g for 5 min at 4°C and resuspended in DPBS (GIBCO) supplemented with 5% FBS, and then stained with the following antibodies: EpCAM-BV785 (BioLegend #118245), CD45-APC (BioLegend, 103112), CD31-APC/Cy7 (BioLegend, 102534), I-A/I-E - AlexaFluor488 (MHC-II) (BioLegend, 107616) and 4’, 6-diamidino-2-phenylindole (DAPI) (Life Technologies, 62248) was used to eliminate dead cells. The cells were then sorted on the BD FACS Aria III system (BD Bioscience) and the data was analyzed using FlowJo software (BD Bioscience).

### Human and mouse cell cultures

Human pulmonary alveolar epithelial cells (HPEpiC) (ScienCell Research Laboratories, 3200) were supplied at passage 1 and were used directly for experiments. Briefly, HPEpiC were seeded at a density of 1.5e4 cells per well in 96-well CellCarrier plates (PerkinElmer) or 3e5 cells per well in 6 well tissue culture plates (Corning) and cultured in AEpiCM (ScienCell Research Laboratories). The cells were cultured for 24 hours prior to being treated with: Human IL11 (UniProtKB:P20809, GenScript), human TGFβ1 (PHP143B, Bio-Rad), anti-IL11 antibody (X203, Aldevron), IgG antibody (IIE10, Aldevron) or U0126 (Cell Signaling Technology, 9903). For mouse AT2 cell cultures, *tdTomato* positive (*tdT*^+^) cells were isolated from *Sftpc-tdT* mice lungs by FACS sorting for CD45^-^ CD31^-^ EpCAM^+^ *tdT*^+^ cells. The FACS sorted cells were then seeded at a density of 2e4 cells per well in rat tail collagen (Invitrogen, A1048301) coated 96-well CellCarrier plates (PerkinElmer) and cultured in DMEM supplemented with 10% FBS. The cells were allowed to adhere for 24 hours prior to treatment with mouse IL11 (UniProtKB: P47873, GenScript) or mouse TGFβ1 (R&D Systems, 7666-MB).

### Operetta high-content immunofluorescence imaging and analysis

Immunofluorescence imaging and quantification of human pulmonary alveolar epithelial cells (HPAEpiC, (ScienCell Research Laboratories) were performed on the Operetta High Content Imaging System (PerkinElmer) as previously described (*22*). The cells were first fixed in 4% paraformaldehyde (Thermo Fisher Scientific) and permeabilized with 0.1% Triton X-100 in phosphate-buffered saline (PBS). Cell proliferation was monitored using EdU-AlexaFluor488 followed by using the Click-iT EdU Labelling kit (Thermo Fisher Scientific, C10350) according to manufacturer’s protocol. The cells were then incubated with the following primary antibodies: KRT8 (Merck Millipore, MABT329, 1:100), Collagen I (Abcam, ab260043, 1:100), fibronectin (Abcam, ab2413, 1:100), SNAIL (Invitrogen, PA5-85493, 1:100), IL11RA (Abcam, ab125015, 1:100), gp130 (Thermo Fisher Scientific, PA5-28932, 1:100), IL6RA (Thermo Fisher Scientific, MA1-80456, 1:100), SFTPC (Abcam, ab211326, 1:100) or AGER (R&D Systems, MAB1179, 1:100) and visualized using Alexa Flour 488-conjugated secondary antibodies. Plates were scanned and images were collected with the Operetta high-content imaging system (PerkinElmer). Each treatment condition was run in duplicate wells, and 14 fixed fields were imaged and analyzed per treatment group. The percentage of proliferating cells (EdU^+ve^ cells) and SNAIL^+^ cells were quantified using the Harmony software version 3.5.2 (PerkinElmer). Quantification of immunostaining intensity was performed using the Columbus software (version 2.7.2, PerkinElmer), and fluorescence intensity was normalized to the cells area.

### RNA-seq

Total RNA was isolated from HPAEpiC with or without TGFβ1 or IL11 stimulation using RNeasy columns (Qiagen). RNA was quantified using Qubit™ RNA Broad Range Assay Kit (Life Technologies) and assessed for degradation based on RNA Quality Score (RQS) using the RNA Assay and DNA 5K/RNA/CZE HT Chip on a LabChip GX Touch HT Nucleic Acid Analyzer (PerkinElmer). TruSeq Stranded mRNA Library Prep kit (Illumina) was used to assess transcript abundance following standard instructions from the manufacturer. Briefly, poly(A)+ RNA was purified from 1ug of total RNA with RQS > 9, fragmented, and used for cDNA conversion, followed by 3’ adenylation, adaptor ligation, and PCR amplification. The final libraries were quantified using Qubit™ DNA Broad Range Assay Kit (Life Technologies) according to the manufacturer’s guide. The quality and average fragment size of the final libraries were determined using DNA 1K/12K/Hi Sensitivity Assay LabChip and DNA High Sensitivity Reagent Kit (PerkinElmer). Libraries with 16 unique dual indexes were pooled and sequenced on a NextSeq 500 benchtop sequencer (Illumina) using NextSeq 500 High Output v2 kit and 75-bp paired-end sequencing chemistry.

### RNA-seq analysis

Libraries were demultiplexed using bcl2fastq v2.20.0.422 with the --no-lane-splitting option. Adapter sequences were then trimmed using trimmomatic v0.36 (*55*) in paired end mode with the options MAXINFO:35:0.5 MINLEN:35. Trimmed reads were aligned to the Homo sapiens GRCh38 using STAR v.2.7.9a (*56*) with the options --outFilterType BySJout -- outFilterMultimapNmax 20 --alignSJoverhangMin 8 --alignSJDBoverhangMin 1 -- outFilterMismatchNmax 999 --alignIntronMin 20 --alignIntronMax 1000000 --alignMatesGapMax 1000000 in paired end, single pass mode. Only unique alignments were retained for counting. Counts were calculated at the gene level using the FeatureCounts module from subread v.2.0.3 (*57*), with the options -O -s 2 -J -T 8 -p -R CORE -G. The Ensembl release 104 Homo sapiens GRCh38 GTF was used as annotation to prepare STAR indexes and for FeatureCounts. Differential expression analyses were performed in R v4.2.0 using the Bioconductor package DESeq2 v1.36.0 (*58*), using the Wald test for comparisons. For sample groups, the design for the model was specified as ∼ stimulus (IL11/TGFβ1/baseline) + source (commercial tube 1-4), to account for the confounding effect of different batches of cells.

### Lung histology and immunohistochemistry

Mouse lung tissue (left lobes) were fixed in 10% formalin for 16-20 hr, dehydrated and embedded in paraffin and sectioned (7 μm) for Masson’s trichrome staining as described previously (*22*). Histological analysis for fibrosis was performed blinded to genotype and treatment exposure as previously described (*22*). For immunostaining, the lungs were fixed in 4% paraformaldehyde at 4°C for 6-16 hrs followed by serial 15% to 30% sucrose in PBS dehydration for 48 hours. The tissues were then embedded in OCT compound prior to sectioning (10 μm). The sections were incubated overnight at 4°C with the following primary antibodies: KRT8 (Merck Millipore, MABT329, 1:100), p-ERK (Cell Signaling Technology, 4370, 1:100), Ki67 (Abcam, ab16667, 1:100), SFTPC (Abcam, ab211326, 1:100), GFP (Abcam, ab290 / ab6673, 1:100), AGER (R&D Systems, MAB1179, 1:200), Podoplanin (R&D Systems, AF3244, 1:200), IL11 (Invitrogen, PAS-95982, 1:100), CD45 (Proteintech, 20103-1-AP, 1:50), PDGFRA (R&D Systems, AF1062, 1:100). Alexa Fluor-conjugated secondary antibodies (Invitrogen, 1:500) were incubated at room temperature for 60 min. Nuclei were stained with DAPI (Invitrogen, 1:1000). Images were captured using the Leica DMi8 microscope (Leica Microsystems) with a 20X or 40X objective. The cells were counted based on positive staining for immunohistochemistry markers and DAPI using Fiji software. Five to ten non-overlapping images for each unique marker were analyzed per mouse lung and the mean values per mouse were presented.

### Colorimetric assays

Detection of secreted IL11 into the supernatant of HPAEpiC cultures was performed using the human IL-11 ELISA kit (R&D systems, D1100) according to manufacturers’ instructions. Detection of SFTPD in mouse serum was performed using the mouse SP-D ELISA kit (ab240683) according to the manufacturer’s instructions. Total lung hydroxyproline content of the right lobes of mice were measured using the Quickzyme Total Collagen assay kit (Quickzyme Biosciences) as previously described (*22*).

### Statistical analysis

Statistical analyses were performed using Graphpad Prism (v9). Analyses were performed using two-tailed Student’s *t*-test, or one-way ANOVA as indicated in the figure legends. For comparisons between multiple treatment groups, *P* values were corrected for multiple testing using Sidak’s test or Tukey’s test. *P* values <0.05 were considered statistically significant.

## Acknowledgements

We thank J.Dong, D.Yeong and B.George for their technical and administrative support.

## Author contributions

B.N. and S.A.C conceived and designed the study. K.Y.H. and C.J.P. performed computational analyses. B.N. and S.V. performed in vitro experiments. B.N., W.-W.L. and F.K. performed in vivo studies and histological evaluations. C.J.P. and H.A.A. performed RNA-seq experiments. B.N., E.P. and S.A.C. supervised the study. B.N. and S.A.C. wrote the manuscript with input from all other authors.

## Data and materials availability

All data associated with this study are presented in the paper or in the Supplementary Materials. RNA sequencing data generated for this study are currently being uploaded onto GEO.

## Supplementary Figures

**Fig S1.**
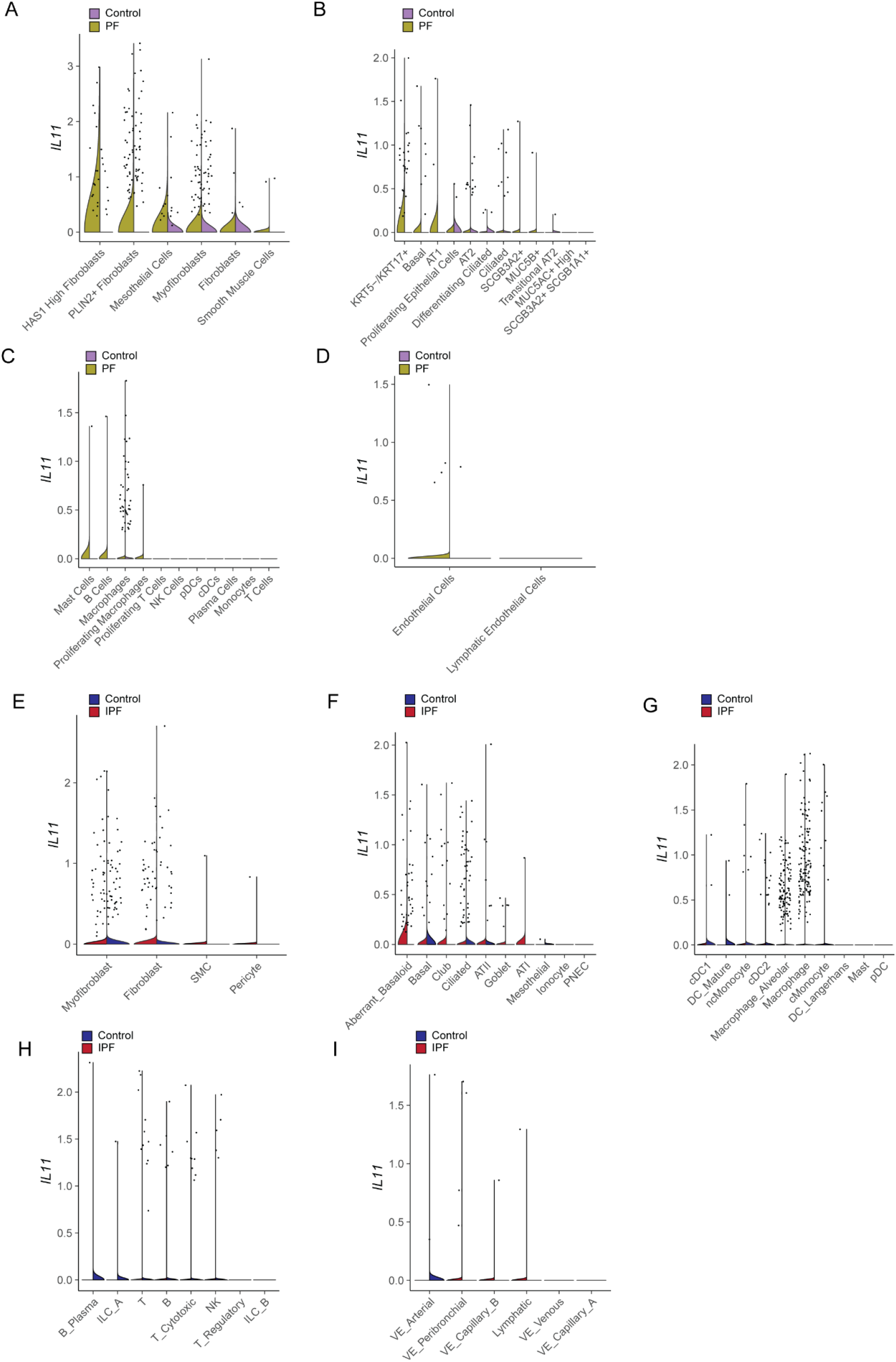
*IL11* is elevated across mesenchymal and epithelial cell subsets in human pulmonary fibrosis. Violin plot of *IL11* expression in individual cell types in scRNA-seq data from control and PF samples in (**A-D**) Habermann et. al. (GSE135893) and (**E-I**) Adams et. al. (GSE136831) datasets. Data are further grouped by PF or control in mesenchymal, epithelial, immune (myeloid or lymphoid) or endothelial cell types.

**Fig. S2.**
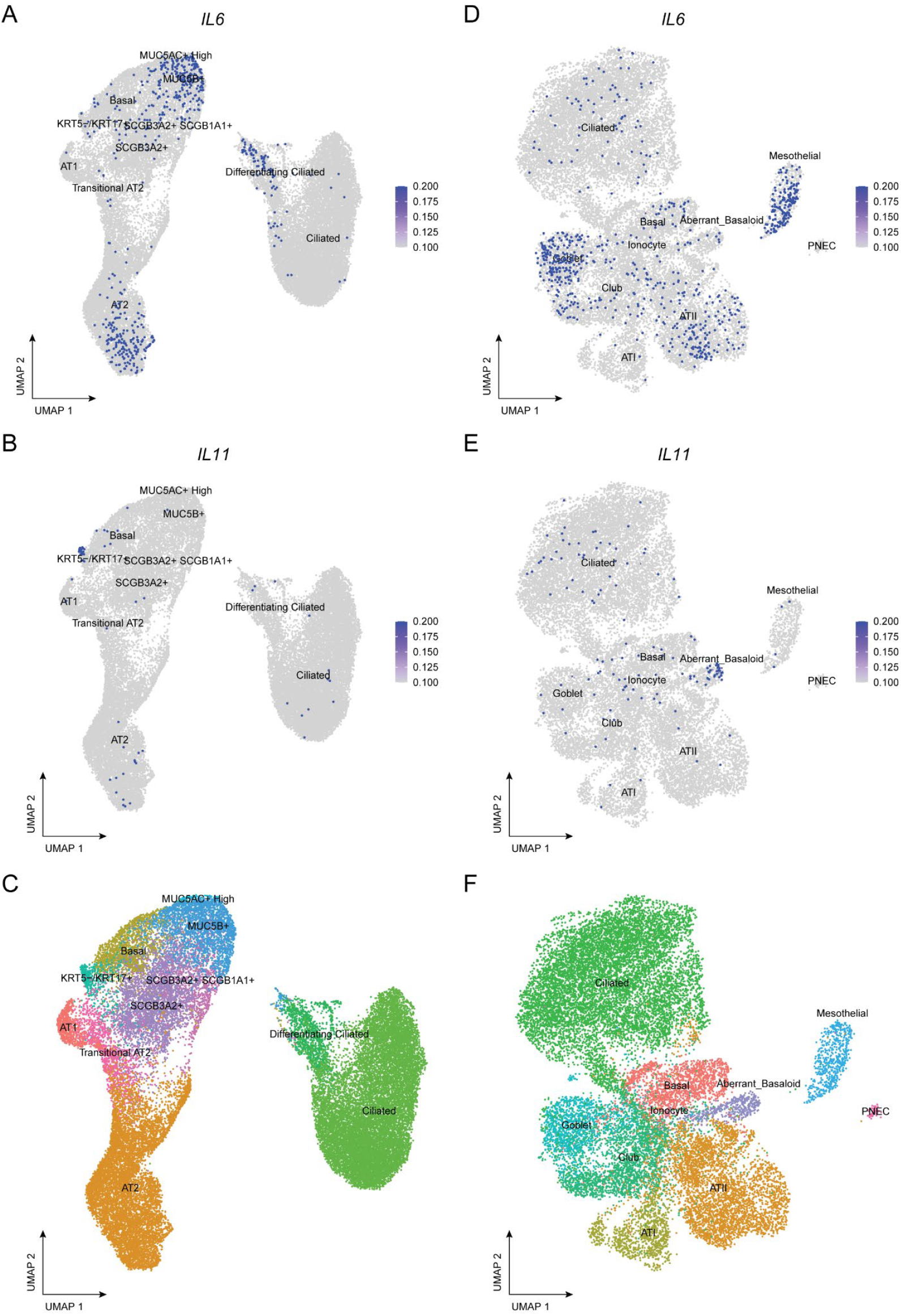
*IL6* and *IL11* expression in various epithelial cell types in the human lung. UMAP visualization of *IL6* or *IL11* expressing single cells in scRNA-seq data from control and PF samples in the (**A-C**) Habermann et. al. (GSE135893) and (**D-F**) Adams et. al. (GSE136831) dataset. *IL6 or IL11* expressing cells are colored in dark blue in A, B, D, E and colored dots indicate different cell clusters in C and F.

**Fig. S3.**
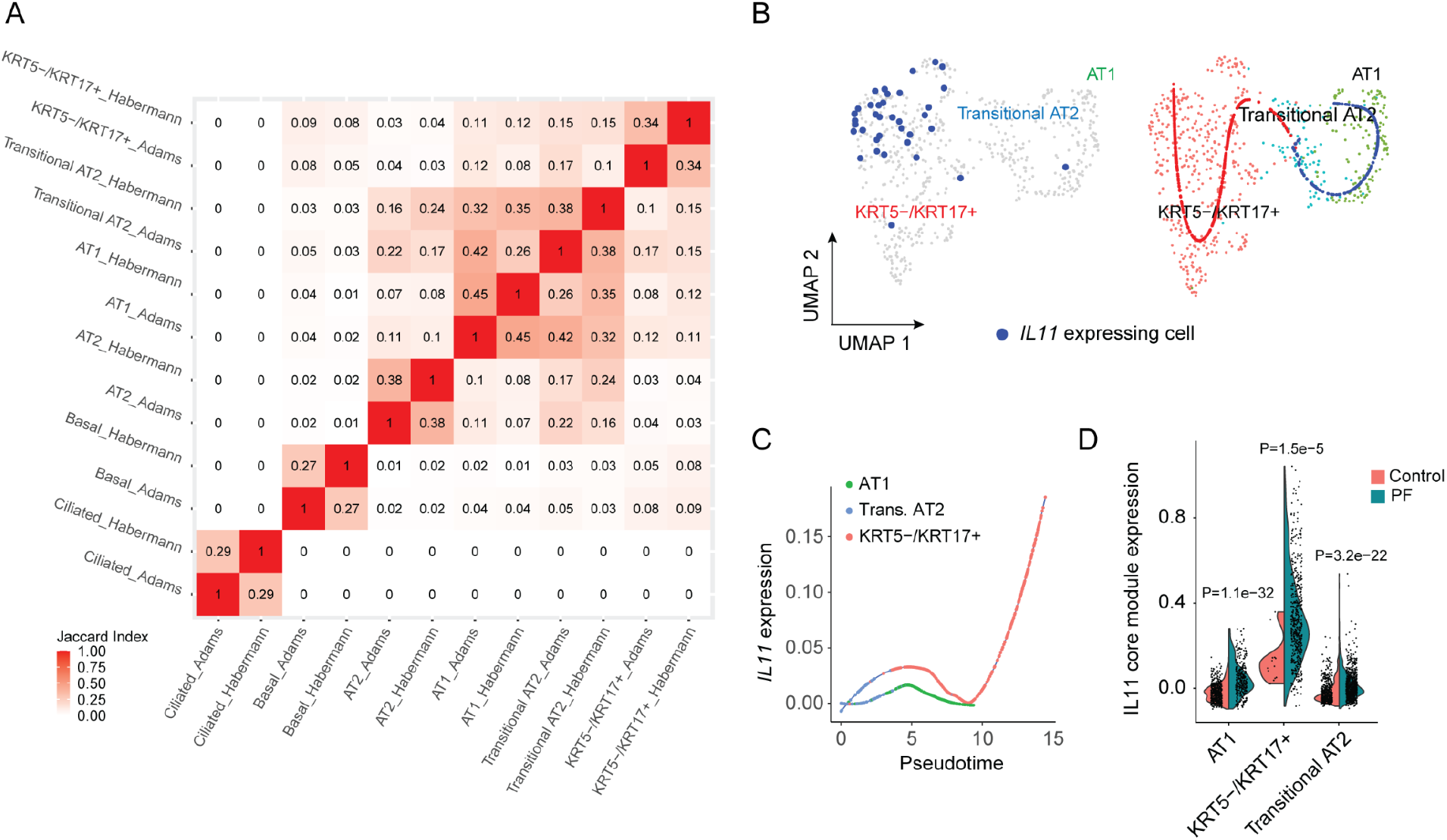
*IL11* is expressed by aberrant basaloid cells in human pulmonary fibrosis. (**A**) Heatmap showing the transcriptional similarities between selected epithelial cell types from Habermann et. al., (GSE135893) and Adams et. al. (GSE136831) datasets as assessed by the Jaccard index. (**B**) UMAP visualization of *IL11* expressing single cells in the Adams et. al. dataset. *IL11* expressing cells are colored in dark blue (left panel) and colored dots indicate cell type clustering (right panel). Cell labels were assigned using the classification from Habermann et. al. by Seurat’s FindTransfer Algorithm (see Methods). Blue line indicates differentiation trajectory from transitional AT2 to AT1 cells; red line indicates differentiation trajectory from transitional AT2 to KRT5^-^/KRT17^+^. Data are composed of PF samples in the Adams et. al. dataset. (**C**) Expression of *IL11* in the pseudotime trajectory from transitional AT2 to KRT5^-^ /KRT17^+^ versus from transitional AT2 to AT1 cells. Data are composed from PF samples in the Adams et. al. dataset. (**D**) Expression of *IL11* co-expression module in transitional AT2, KRT5^-^/KRT17^+^ and AT1 cells from control versus PF in combined Habermann et. al. and Adams et. al. datasets.

**Fig. S4.**
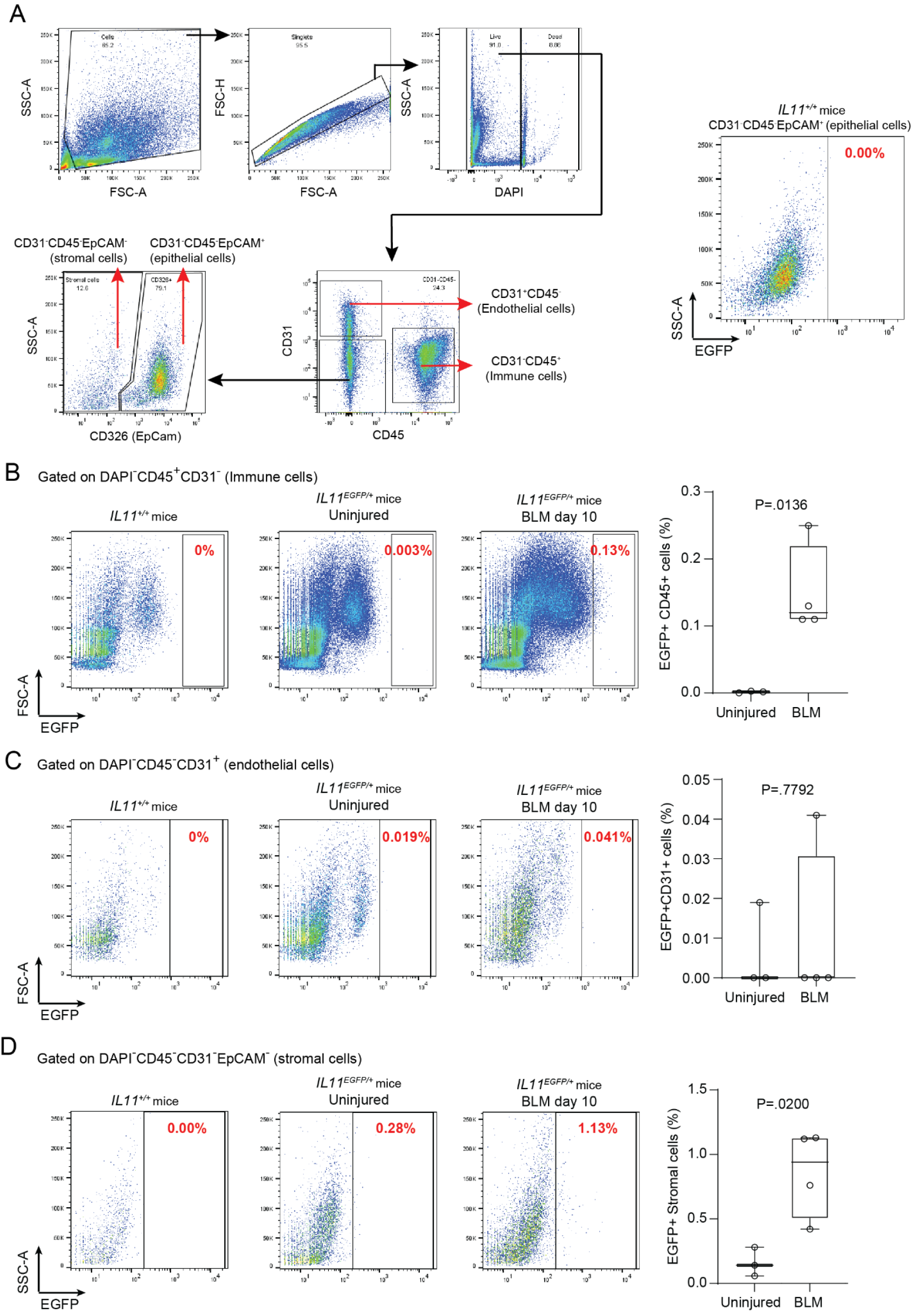
Flow cytometry gating of EGFP^+^ cells from *IL11*^*EGFP*^ reporter mice lungs. (**A**) Representative gating for flow cytometry analysis of lung stromal and epithelial cells isolated from *IL11*^*EGFP*^ reporter mice based on antibody staining for CD31, CD45 or CD326 (EpCAM). Rightmost panel indicates the predetermined threshold for IL11^EGFP+^ expression based on the EGFP signal in CD31^-^ CD45^-^ CD326^+^ epithelial cells from *IL11*^*+/+*^ littermate control mice. Percentages of (**B**) EGFP^+^ immune cells (CD31^-^ CD45^+^), (**C**) EGFP^+^ endothelial cells (CD31^+^CD45^-^) and (**D**) EGFP^+^ stromal cells (CD31^-^CD45^-^EpCAM^-^) in uninjured and bleomycin (BLM)-treated *IL11*^*EGFP*^ reporter mice based on the EGFP signal in the respective cell types in *IL11*^*+/+*^ control. *P* value determined by two-tailed Student’s *t-*test, n = 4 mice. Data are represented as median and whiskers extend from minimum to maximum values.

**Fig. S5.**
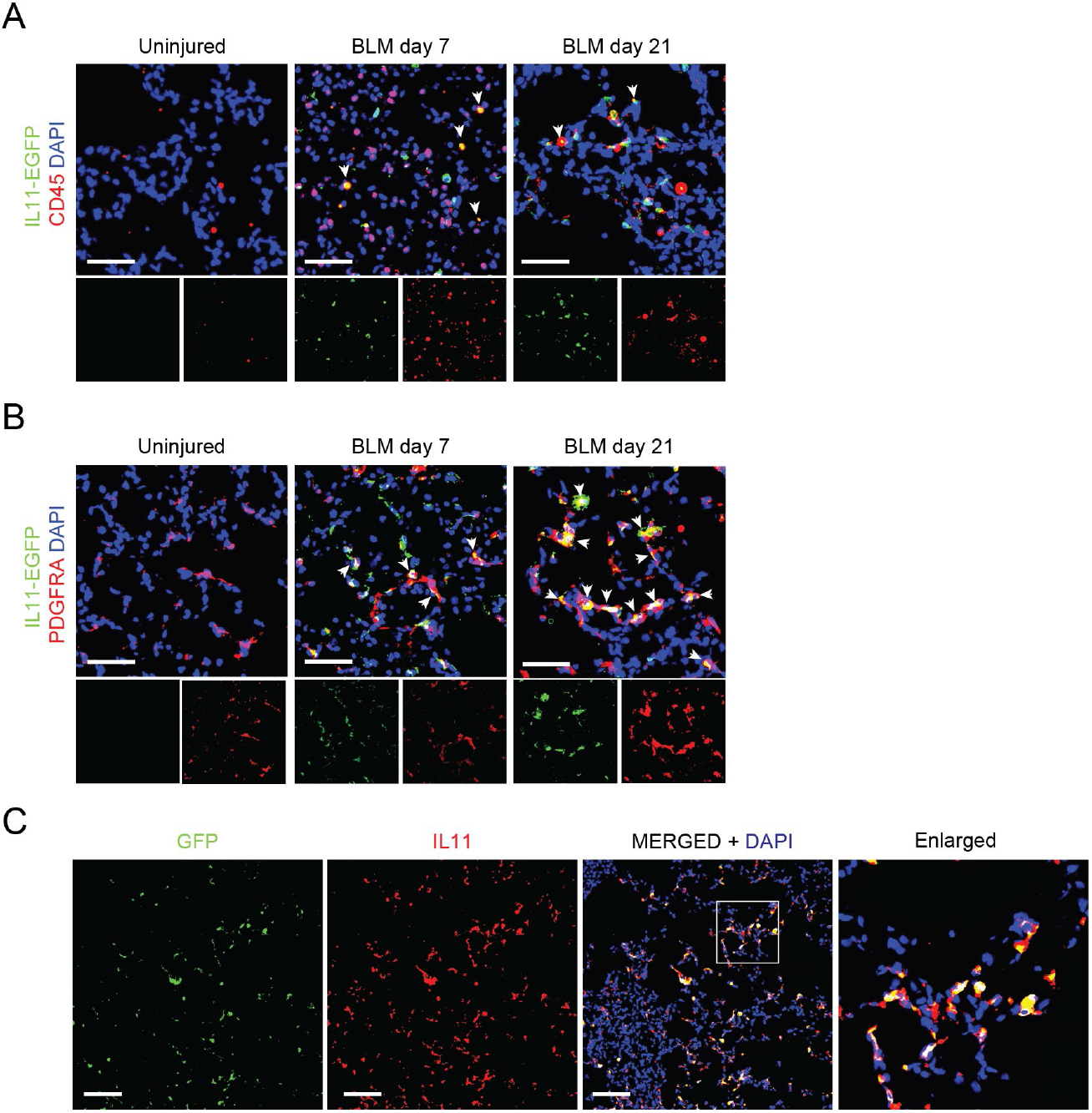
IL11 is upregulated in lung fibroblasts and CD45^+^ cells after bleomycin-induced lung injury in mice. Immunostaining for GFP and (**A**) CD45 or (**B**) PDGFRA in the lungs of *IL11*^*EGFP*^ reporter mice post-BLM injury. White arrowheads indicate marker positive IL11^EGFP+^ cells. Scale bars: 50 μm. (**C**) Immunostaining of lungs from *IL11*^*EGFP*^ reporter mice post-BLM injury with anti-GFP and anti-IL11 antibodies, showing considerable overlap of GFP and IL11 signals. Scale bars: 100 μm.

**Fig. S6.**
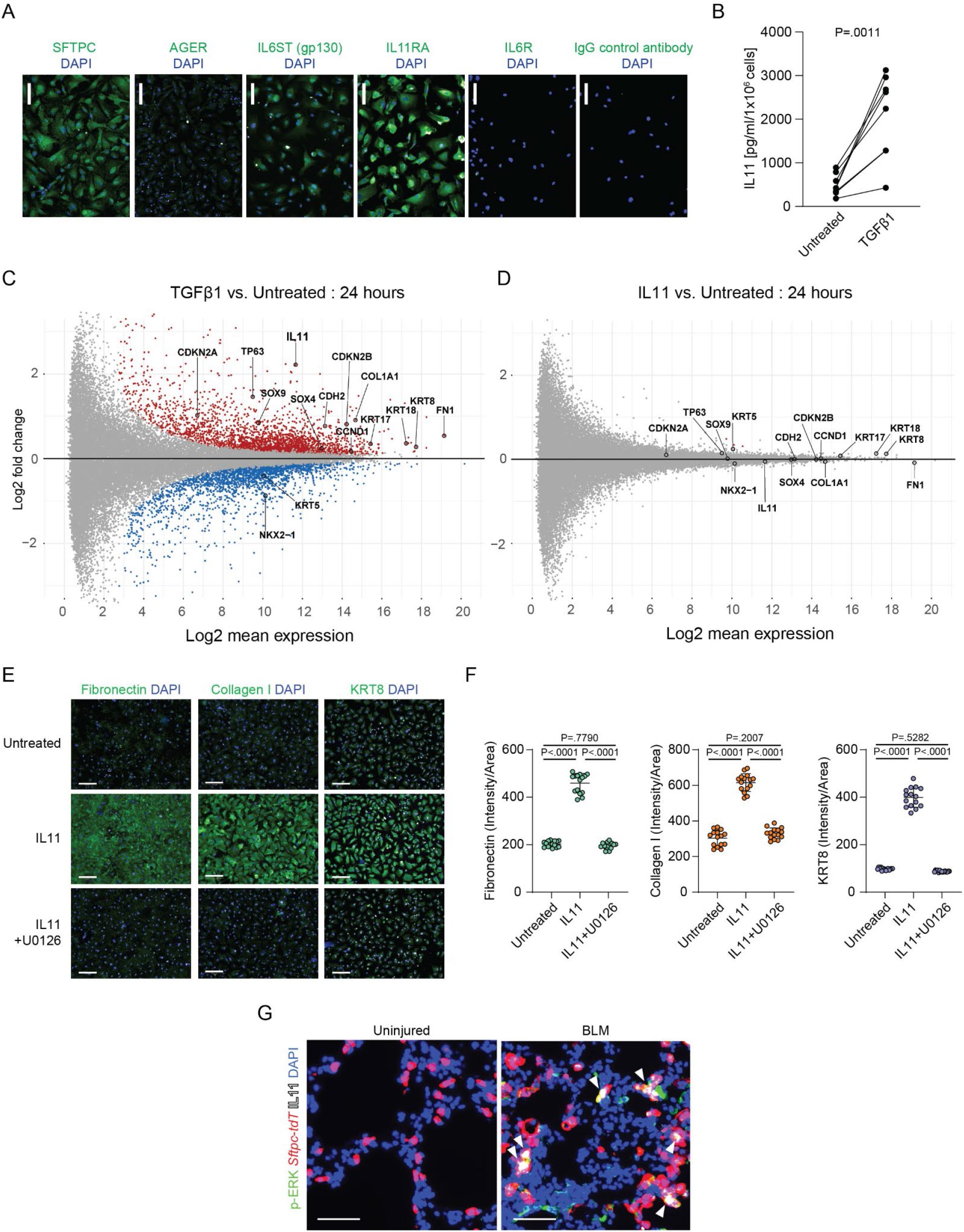
IL11-dependent ERK signaling promotes cytopathic features in AT2 cells *in vitro*. (**A**) Representative images of immunostaining for SFTPC, AGER, IL6ST (gp130), IL11RA and IL6R in primary human pulmonary alveolar epithelial cells. Scale bars: 100 μm. (**B**) ELISA-based quantification of IL11 in culture supernatant of TGFβ1-treated HPEpiC (5 ng/ml, 24 hours, n = 6). *P* value determined by two-tailed paired *t*-test. (**C-D**) RNA-seq analysis of TGFβ1- or IL11-treated human pulmonary alveolar epithelial cells (5 ng/ml, 24 hours). Red and blue dots indicate significantly differentially upregulated or downregulated genes respectively (FDR < 0.05). (**E**) Representative images of immunostaining of Fibronectin, Collagen I and KRT8 in HPEpiC treated with IL11 (5 ng/ml) in the presence of MEK inhibitor U0126 (10 μM) for 24 hours. Scale bars: 100 μm. (**F**) Quantification of immunostaining in panel **E**. Data are represented as mean ± s.d. *P* values were determined by one-way ANOVA (Tukey’s test). (**G**) Images of immunostaining of IL11 and p-ERK in lung sections from *Sftpc-tdT* mice after BLM-induced lung injury. Scale bars: 50 μm. White arrowheads indicate p-ERK^+^ IL11^+^ *tdT*^+^ cells.

**Fig. S7.**
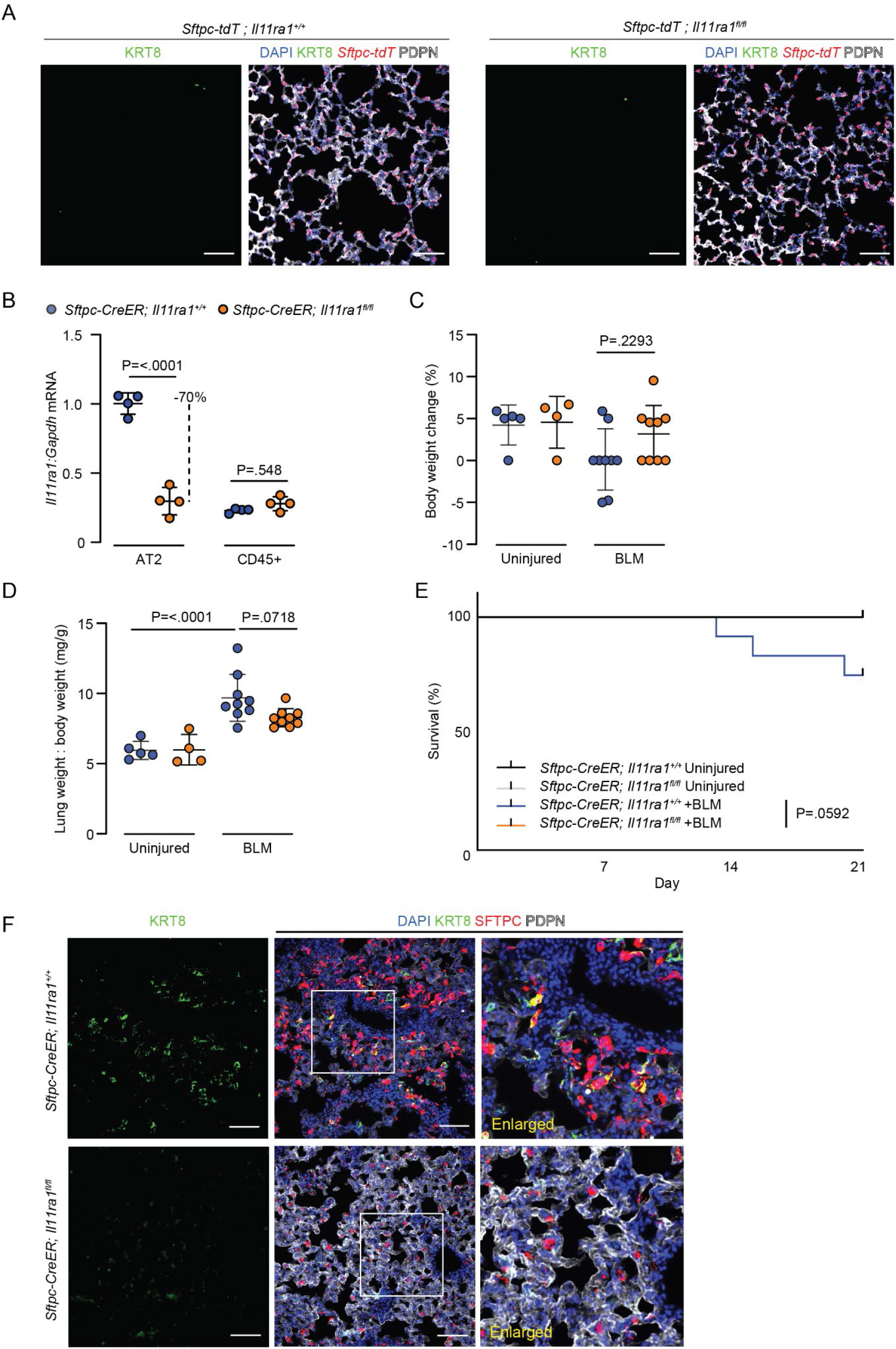
Characterization of AT2 cell-specific *Il11ra1*-deleted mice after bleomycin injury. (**A**) Immunostaining for KRT8 and PDPN in lungs of uninjured *Sftpc-tdT; Il11ra1*^*fl/fl*^ *or Il11ra1*^*+/+*^ mice. Scale bars: 100 μm. (**B**) qPCR analysis of *Il11ra1* expression in AT2 cells (CD45^-^ CD31^-^ EpCAM^+^ MHCII^+^) and CD45^+^ immune cells isolated from the lungs of *Sftpc-CreER; Il11ra1*^*fl/fl*^ and *Sftpc-CreER; Il11ra1*^*+/+*^ mice. (**C**) Percentage body weight change post-BLM injury (day 21 versus day 0), (**D**) Lung weight to body weight indices and (**E**) survival analysis of *Sftpc-CreER; Il11ra1*^*fl/fl*^ *or Il11ra1*^*+/+*^ mice 21 days post-BLM treatment. (**F**) Images of immunostaining for KRT8, SFTPC and PDPN in lung sections from *Sftpc-CreER; Il11ra1*^*fl/fl*^ *or Il11ra1*^*+/+*^ mice 21 days post-BLM treatment. Scale bars: 100 μm. Data are represented as mean ± s.d. *P* values were determined by two-tailed Student’s *t*-test in panel **A** and one-way ANOVA (Tukey’s test) in panel **B** to **D** and by Log-rank (Mantel-Cox) test in panel **E**.

**Fig. S8.**
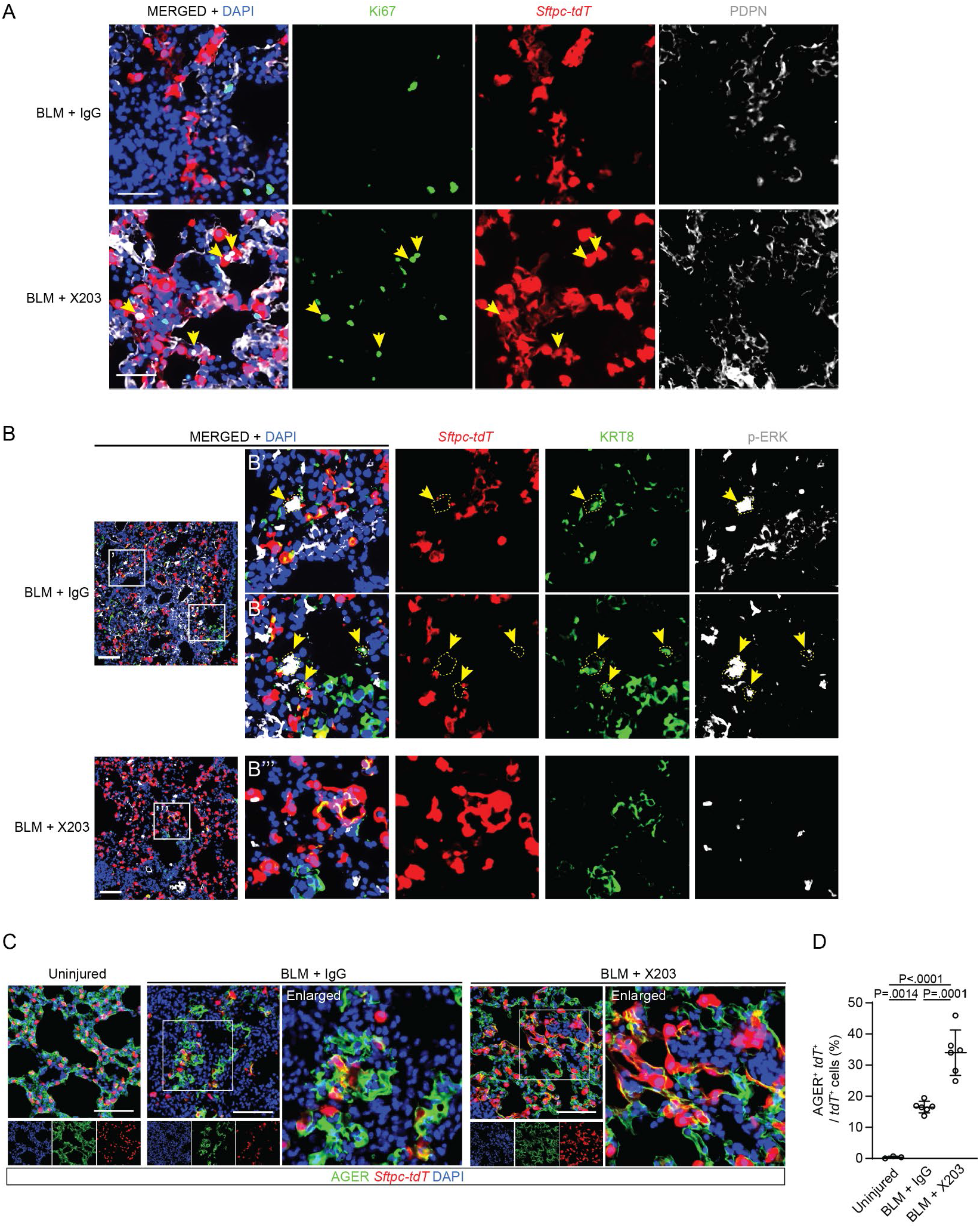
Pharmacological inhibition of IL11 promotes AT2 cell proliferation and reduced p-ERK+ KRT8+ cells after bleomycin-induced lung injury. (**A**) Representative images of immunostaining for Ki67 and PDPN or (**B**) KRT8 and p-ERK in lung sections from BLM-injured *Sftpc-CreER; R26-tdT* mice treated with X203 or IgG antibodies. Scale bars: 50 μm in A and 100 μm in B. Yellow arrowheads indicate Ki67^+^ *tdT*^+^ cells in panel A or p-ERK^+^ KRT8^+^ *tdT*^+^ cells in panel B. (**C**) Representative images of immunostaining for AGER and (**D**) the ratio of AGER^+^ *tdT*^+^ to *tdT*^+^ cells in lung sections from BLM-injured *Sftpc-CreER; R26-tdT* mice treated with X203 or IgG antibodies. Scale bars: 100 μm. Data are represented as mean ± s.d. *P* values were determined by one way ANOVA (Tukey’s test), n = 3 - 6 mice / group.

